# The spatial chronnectome reveals a dynamic interplay between functional segregation and integration

**DOI:** 10.1101/427450

**Authors:** A. Iraji, T.P. DeRamus, N. Lewis, M. Yaesoubi, J.M. Stephen, E. Erhardt, A. Belger, J.M. Ford, S. McEwen, D.H. Mathalon, B.A. Mueller, G.D. Pearlson, S.G. Potkin, A. Preda, J.A. Turner, J.G. Vaidya, T.G.M. van Erp, V.D. Calhoun

## Abstract

The brain is highly dynamic, reorganizing its activity at different interacting spatial and temporal scales including variation within and between brain networks. The chronnectome is a model of the brain in which nodal activity and connectivity patterns are changing in fundamental and recurring ways through time. Most previous work has assumed fixed spatial nodes/networks, ignoring the possibility that spatial nodes or networks may vary in time, particularly at the level of the voxel. Here, we introduce an approach allowing for a spatially fluid chronnectome (called the spatial chronnectome for clarity), which focuses on the variation in spatiotemporal coupling at the voxel level within each network. We identify a novel set of spatially dynamic features which can be obtained and evaluated under different conditions. Results reveal transient spatially fluid interactions between intra- and inter-network relationships in which brain networks transiently merge and then separate again, emphasizing the dynamic interplay between segregation and integration. We also show that brain networks exhibit distinct spatial patterns with unique temporal characteristics, potentially explaining a broad spectrum of inconsistencies in previous studies which assumed static networks. Moreover, we show for the first time that anticorrelative connections to the default mode network, are transient as opposed to constant across the entire scan. Preliminary assessments of the approach using a multi-site dataset collected from 160 healthy subjects and 149 patients with schizophrenia (SZ) revealed the ability of the approach to obtain new information and nuanced alterations of brain networks that remain undetected during static analysis. For example, patients with SZ display transient decreases in voxel-wise network coupling including within visual and auditory networks that are not detectable in a spatially static analysis. Our approach also enabled calculation of a novel parameter, the intra-domain coupling variability which was higher within patients with SZ. The significant association between spatiotemporal uniformity and attention/vigilance cognitive domain highlights the cognitive relevance of the spatial chronnectome. In summary, the spatial chronnectome represents a new direction of research enabling the study of functional networks that are transient at the voxel level and identification of mechanisms for within and between-subject spatial variability to study functional brain homeostasis.

## Introduction

Neuroimaging modalities provide unique opportunities to model and investigate brain functional connectivity at a large/macro scale. One key finding is a set of replicable large-scale functional brain networks (Biswal et al., 2010; Buckner et al., 2009; Franco et al., 2009; Guo et al., 2012; Shehzad et al., 2009; Zuo et al., 2010). Brain networks, and groups of temporally coherent activity within networks called functional domains, have been studied and validated using various analytical approaches (Calhoun et al., 2008; Smith et al., 2009; van den Heuvel et al., 2009; Van Dijk et al., 2010; Yeo et al., 2011). Of these approaches, independent components analysis (ICA) enables simultaneous extraction of both the spatial patterns of functional domains and their temporal activity patterns. Studies of brain networks and functional domains demonstrate alterations in their spatial/temporal patterns under different physiological and psychological conditions, as well as disease (Arbabshirani et al., 2017; Garrity et al., 2007; Greicius, 2008; Iraji et al., 2015; Menon, 2011; Seeley et al., 2009; Sorg et al., 2007). However, these studies hold a common assumption that each brain network is comprised of a fixed set of brain regions with a static pattern of activity over time. This is an oversimplification, as the brain is highly dynamic, with variations in associated regions and spatial patterns of brain functional organizations including brain networks (Calhoun et al., 2014). As such, many recent studies have demonstrated the ability of fMRI to capture time-varying brain connectivity (Calhoun et al., 2014; Hutchison et al., 2013; Preti et al., 2017). For instance, studying whole-brain dynamic connectivity demonstrates variations in temporal coupling, both within and between functional domains (Allen et al., 2014; Damaraju et al., 2014). Examining temporal coupling between brain regions reveals strong correlations between regions known to be part of one network with regions of other networks at particular moments in time. This suggests “isolated” brain networks are only transiently isolated. Additionally, despite recent developments in detecting the dynamic behavior of brain activity using fMRI, spatiotemporal variations of brain networks have been underappreciated. Previous time-varying studies have focused on either 1) dynamic temporal coupling among fixed spatial nodes/networks, which ignore the importance of spatial variations (Allen et al., 2014; Barttfeld et al., 2015; Chen et al., 2016; Damaraju et al., 2014; Hutchison et al., 2013; Leonardi et al., 2013; Sakoglu et al., 2010; Shine et al., 2016; Yaesoubi et al., 2018) or 2) the dominant spatial pattern at any given time using a single region or whole brain time courses without capturing the spatiotemporal variations within and between functional organizations (Karahanoglu and Van De Ville, 2015; Liu and Duyn, 2013; Preti and Van De Ville, 2017; Tagliazucchi et al., 2012; Trapp et al., 2018; Vidaurre et al., 2017). Kiviniemi and his colleagues presented work highlighting spatial variation in the default mode network using sliding-window ICA (Kiviniemi et al., 2011). Other work investigated fluctuation in spatial couplings between spatial components derived from independent vector analysis (Ma et al., 2014). While these present intriguing early evidence, to date, there has not been an approach that has evaluated the spatial fluidity within and between functional brain networks.

The chronnectome is a model of the brain in which nodal activity and connectivity patterns are changing in fundamental and recurring ways through time (Calhoun et al., 2014). Here, we introduce an approach allowing for a spatially fluid chronnectome (called the *spatial* chronnectome) which advances current analytical methods by providing, novel, time-varying information of individual brain networks at the voxel level. Encapsulating transient voxel-wise network coupling allows researchers to capture the spatially fluid behavior of brain networks. To investigate the spatiotemporal variations of brain networks, we calculated the relationship (temporal correlation) of each individual brain network with every voxel of the brain. Because the time course of each brain network obtained from ICA represents its temporal activity, its temporal correlation with each brain voxel provides information about the involvement of the voxel to the brain network. Therefore, the spatiotemporal variations of a brain network were encapsulated through measuring the coupling (temporal correlation) between every brain voxel and the given brain network at different moments using a sliding-window approach. Preliminary assessments demonstrate the ability of the approach to obtain new information of brain function. A new set of spatially relevant features can be calculated and used to study brain function. For instance, we introduce a metric called the “spatiotemporal transition matrix” to summarize the spatiotemporal information of each brain network. We also characterize distinct spatial states within each brain network with unique temporal characteristics which are highly replicable. Using spatial states, the dynamic properties of each brain network can also be assessed by calculating temporally derived indices such as mean dwell time or fraction time for individual brain networks. It is worth mentioning that the spatial states of brain networks can also be related to interdigitated networks previously only observed in single subject analysis (Braga and Buckner, 2017; Laumann et al., 2015). The approach was further used to evaluate spatial chronnectome properties in patients with schizophrenia (SZ). We hypothesized spatial chronnectome will allow us to detect nuanced alterations in brain networks in patients with SZ, which would be remained undetected during static analyses. The statistical comparison reveals that transient decreases in voxel-wise networks couplings are more pronounced than the reduction in static functional connectivity. Furthermore, a spatial chronnectome analysis detects alterations in brain networks that are not identified when used a spatially static analysis. Using variation-based analysis, we demonstrate, for the first time, higher coupling variability and different spatiotemporal transition patterns across various brain networks. We conclude that the study of spatiotemporal variations of large-scale brain networks can unveil underappreciated features of the dynamic brain and improve our understanding of cognitive and behavioral neuroscience.

## 2. Method

### 2.1. Outline of our approach

Our approach assessing spatiotemporal variations of individual brain networks includes the following steps (Figure 1(A)):

1. Spatial independent component analysis (sICA; can be either group sICA followed by back-reconstruction or single-subject sICA) applied to obtain large-scale brain networks and their associated time courses (Figure 1(A) *Step 1*). The time course of each brain network obtained from sICA represents the temporal activity of that large-scale brain network. Details can be found in *Section 2.4. Identifying large-scale brain networks: Spatial ICA*.
2. Temporal coupling and sliding-window approaches were employed to capture spatiotemporal variations of the large-scale brain networks. For each brain network and time window, we calculated the correlation between the time course of the brain network and the time courses of every voxel of the brain. The resulting correlation values represent the association (involvement) of all voxels across the brain to the given network at each time window (Figure 1(A) *Step 2*). This results in one dynamic coupling map (dCM) per window for each brain network. This approach, unlike its predecessors such as whole brain dynamic functional network connectivity (dFNC) and co-activation patterns (CAP), provides nuanced information regarding temporal variations of spatial patterns of multiple brain networks simultaneously at the level of the voxel. Details can be found in *Section 2.5. Calculating dynamic coupling maps (dCMs) for each brain network using a sliding-window approach*.
3. Time-varying properties were evaluated for each brain network (Figure 1(A) *Step 3)*. First, the dCMs of each individual network were clustered into a set of distinct spatial patterns called spatial states on which multiple dynamic metrics were calculated and investigated (see *Section 2.6. Calculating the spatial states of each brain network and their dynamic patterns*). Next, the spatial variations of each brain network over time were evaluated by calculating voxel-wise changes in their dCMs (See *Section 2.7. Evaluating the spatial variations of each brain network over time*).

**Figure 1.**
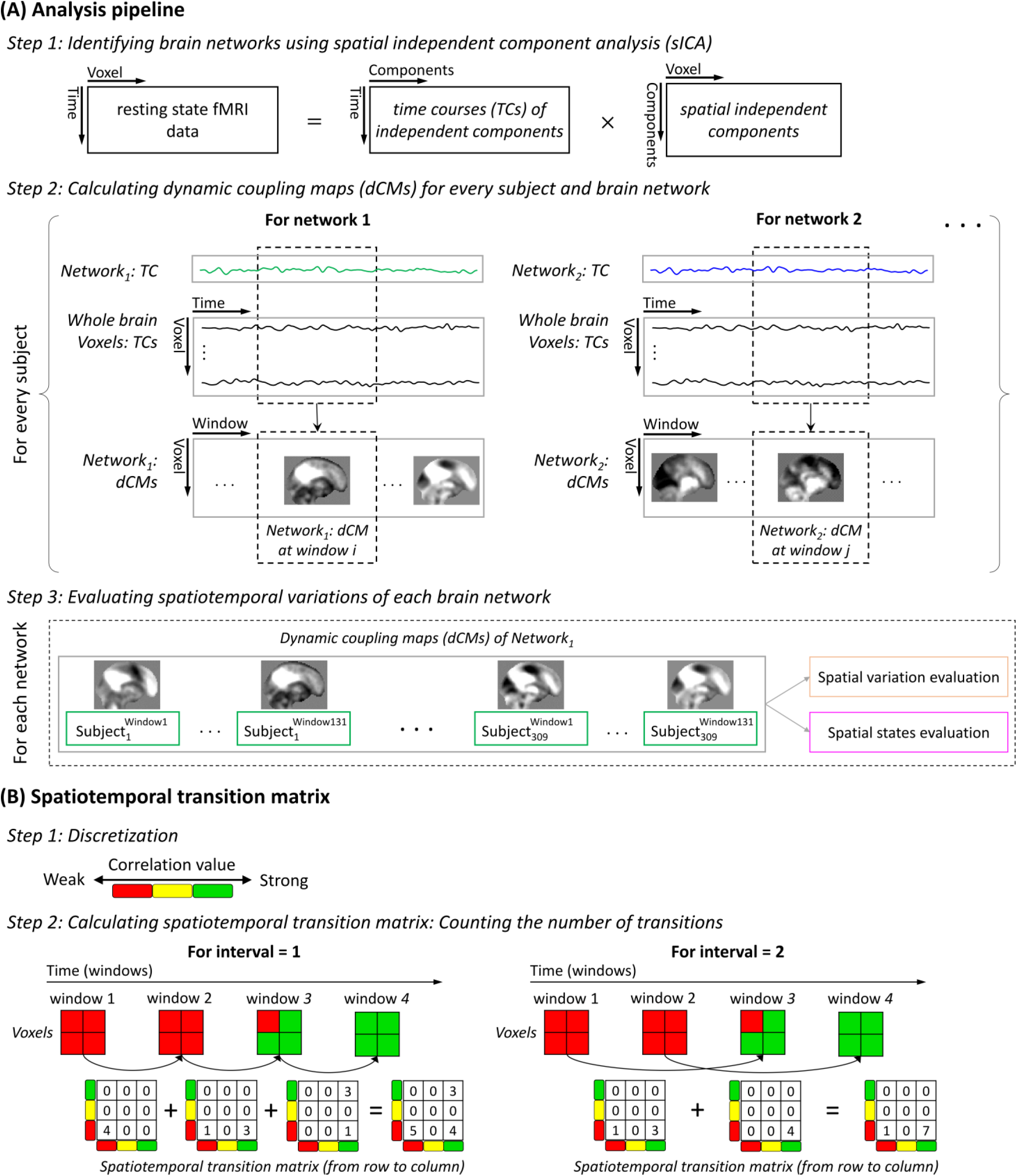
Summary of processing steps. (A) Cartoon of the analysis pipeline. First, spatial independent component analysis (sICA) is applied to obtain brain networks and their associated times courses. ICA consists of several steps including data reduction (principal component analysis), group-level sICA, and subject-level sICA. Following ICA, whole brain dynamic coupling maps (dCMs) of each network for every subject are obtained by calculating windowed-correlation values between the time course of the network and every brain voxel. Finally, time-varying properties of each network are investigated by evaluating spatial variation of dCMs over time and estimating spatial states. Details of the spatiotemporal variation analysis can be found in Sections 2.6 and 2.7. (B) A toy model of calculating the spatiotemporal transition matrix. First, we discretize the correlation value. For visualization purpose, we present three bins here, but 10 bins were used in analysis of the real data. Next, we count the number of transitions from one bin to another between time windows at a given interval.

### 2.2. Data collection

Data collection was performed at 7 imaging sites across the United States, and all analyzed data passed data quality control. All participants were at least 18 years old, and gave written informed consent prior to enrollment. Data were collected from 160 healthy participants, including 46 females and 114 males (average age: 36.71 ± 10.92; range: 19-60 years), and 149 age- and gender-matched patients with SZ, including 36 females and 113 males (average age: 37.95 ± 11.47; range: 18-60 years). The imaging data were collected on a 3-Tesla Siemens Tim Trio scanner for six of the seven sites and on a 3-Tesla General Electric Discovery MR750 scanner at one site. Resting state fMRI (rsfMRI) data was acquired using a standard gradient echo-planar imaging (EPI) sequence with following imaging parameters: repetition time (TR) = 2000 ms, echo time (TE) = 30 ms, flip angle (FA) = 77°, field of view (FOV) = 220 × 220 mm, matrix size = 64 × 64, mm, pixel spacing size = 3.4375 × 3.4375 mm, slice thickness = 4, slice gap = 1 mm, number of excitations (NEX) = 1, and a total of 162 volumes. During rsfMRI scans, participants were instructed to keep their eyes closed and rest quietly without falling asleep. Further details on this dataset can be found in our earlier work (Damaraju et al., 2014).

### 2.3. Preprocessing

The preprocessing was performed primarily using SPM (http://www.fil.ion.ucl.ac.uk/spm/) and AFNI (https://afni.nimh.nih.gov) software packages. The pipeline includes brain extraction, motion correction using the INRIAlign, slice-timing correction using the middle slice as the reference time frame, and despiking using AFNI’s 3dDespike. The data of each subject was subsequently warped to the Montreal Neurological Institute (MNI) template using non-linear registration, resampled to 3 mm^3^ isotropic voxels, and spatially smoothed using a Gaussian kernel with a 6 mm full width at half-maximum (FWHM = 6 mm). Finally, voxel time courses were z-scored (variance normalized) as it has been shown in our analysis to better estimate brain networks relative to other scaling methods for independent component analysis.

### 2.4. Identifying large-scale brain networks: Spatial ICA

Spatial ICA (sICA) was applied to the fMRI data to obtain brain networks (Calhoun and Adali, 2012; Calhoun et al., 2001a). ICA was performed using the GIFT software package (http://mialab.mrn.org/software/gift/) in the following steps similar to our previous work (Iraji et al., 2016): 1) Subject-level principal component analysis (PCA) was applied and the 30 principal components accounting for the maximum variance in each individual dataset were retained. 2) Subject-level principle components were concatenated across time, and group-level spatial PCA was applied. 3) The first 20 principal components with the highest variance were selected as input for the infomax algorithm to estimate the 20 group independent components (Bell and Sejnowski, 1995; Correa et al., 2007; Correa et al., 2005). Infomax ICA algorithm was repeated 100 times on the 20 group-level principal components using ICASSO framework in order to obtain a stable and reliable estimation of independent components (Himberg et al., 2004) and the most representative run was selected for further analysis (Ma et al., 2011). 4) The subject-specific independent components and component time courses were calculated using group information guided ICA (GIG-ICA) (Du and Fan, 2013; Du et al., 2015). 5) The twelve independent components were identified as brain networks based on the spatial and temporal properties and prior knowledge from previous studies.

### 2.5. Calculating dynamic coupling maps (dCMs) for each brain network using a sliding-window approach

The spatiotemporal variations of each brain network can be captured by evaluating its dynamic coupling at a voxel-wise level. For this purpose, we calculated the temporal coupling between the brain network and every voxel of the brain using the sliding-window correlation approach. As a result, we compute dCMs for a given network, which encapsulates the spatiotemporal variations of that brain network at a voxel-wise level. The cleaning procedure was first performed on the time courses of brain networks and every voxel of the brain to reduce noise. The cleaning procedure includes orthogonalizing with respect to estimated subject motion parameters, linear detrending, despiking, and band-pass filtering using a fifth-order Butterworth filter (0.001-0.15 Hz). This cleaning procedure has demonstrated its effectiveness in improving the detection of dynamic patterns in whole-brain dFNC analyses (Allen et al., 2014; Damaraju et al., 2014). We used the tapered window obtained by convolving a rectangle (width = 30 TRs) with a Gaussian (σ = 3 TRs) and the sliding step size of one TR resulting in 131 windows per subject (Allen et al., 2014; Damaraju et al., 2014).

### 2.6. Calculating the spatial states of each brain network and their dynamic patterns

First, we modeled the spatiotemporal fluctuations of each brain network as temporal variations in a set of distinct spatial patterns called spatial states. Clustering approaches can be used to summarize the dCMs of each brain network into a set of spatial states. This allows us to investigate the spatiotemporal variations of the brain network via temporal variations of these distinct spatial states. Here, k-means clustering was employed to detect the spatial states of each brain network. For each brain network, k-means clustering was applied on the 40479 (309 subjects × 131 windows) dCMs of the brain network. K-means clustering was repeated 100 times with different initializations using the k-means++ technique to increase chances of escaping local minima (Arthur and Vassilvitskii, 2007). The correlation distance metric was used to measure the similarity between data points (i.e., the dCMs), as it is more effective in the detection of spatial patterns irrespective of voxel intensities. However, an exploratory analysis using Euclidean distance demonstrated almost identical results. K-means clustering was performed for 3 to 10 clusters, and the spatial maps of the cluster centroids were compared. For each brain network, the maximum number of clusters that provided distinct spatial maps for centroids were selected by visual inspection for further analysis. Thus, each network includes multiple spatial states as defined by the cluster centroids, and the number of spatial states (centroids) can vary between networks. We also compared our numbers of clusters with the estimated number of clusters using the elbow criterion (Damaraju et al., 2014; Yaesoubi et al., 2017). With the exception of the left frontoparietal and subcortical domains, in which the elbow criterion estimates a higher number of clusters than those chosen by visual inspection, the estimated cluster numbers using elbow criterion were the same as the expert selections. Using temporal profiles of the spatial states, various state level and meta-state level dynamic indices can be calculated for each brain network. For example, the mean dwell time (the average of the amount of time that subjects stay in a given state once entering that state) and the fraction time (the proportion of time subjects stay in a given state) can be calculated for each networks. Here, we compared mean dwell time and fraction time between healthy subjects and patients with SZ to show the feasibility of the approach.

### 2.7. Evaluating the spatial variations of each brain network over time

#### Coupling variability map

Coupling variability for each network is defined as the amount of variations in network coupling over time, which is obtained by measuring voxel-wise changes in dCMs using the L1 norm distance (sum of absolute differences). For each voxel, the L1 norm distance represents the variations in a voxel’s membership to a given brain network over time by measuring changes in the sliding-window correlation values between the time courses of a given voxel and the brain network across time. For example, if the correlation values for a given voxel for seven consecutive time windows are c_1_, c_2_, …, and c_7_, the changes in correlation values will be d_1_ = |c_2_-c_1_|, d_2_ = |c_3_-c_2_|, … d_6_ = |c_7_-c_6_|. Therefore, the coupling variability for a given voxel will be equal to d_1_+d_2_+…+d_6_ (Figure S1). The coupling variability map quantifies the overall spatial behavior of a given brain network varying across time. Voxel-wise comparisons were further applied comparing coupling variability maps for each network between healthy subjects and patients with SZ.

#### Spatiotemporal transition matrix

To further evaluate spatiotemporal variations of each brain network, we exploit the gray-level co-occurrence (spatial dependence) matrix method in the field of image processing that is used to extract Haralick textural features from images (Haralick et al., 1973). Figure 1(B) shows a toy model of this approach. First, the network coupling (i.e., voxel-to-network correlation) values are discretized to *n* bins. The spatiotemporal transition matrix is constructed by counting the number of voxels transitioning from one bin to another between time windows at a given interval. We calculate this spatiotemporal transition matrix for interval equal to one TR, (i.e., by counting the number of transitions between every two consecutive spatial maps), and for larger intervals. The maximum interval would be the total number of windows per subject minus 1 (i.e., 130). The spatiotemporal transition matrix was normalized (divided) by the total number of transitions to allow us to compare across different interval values. Several global indices such as contrast, correlation, energy, entropy, and homogeneity can be calculated from spatiotemporal transition matrix to provide summary statistics of spatiotemporal variations of brain networks (Haralick et al., 1973). Here, for example, the energy index was calculated to evaluate spatiotemporal uniformity. The energy index, also known as angular second moment (ASM), is defined as ∑*_i,j_p*(*i,j*)^2^ in which *p*(*i,j*) is the (*i*,*j*)^th^ element of the spatiotemporal transition matrix. The energy index is between zero and one, and smaller energy values represent higher spatiotemporal uniformity. The energy index of each network was compared between healthy subjects and patients with SZ. The energy indices of the networks which show significant differences between the two groups were further analyzed to understand the relationships between the spatiotemporal variations and cognitive scores. For this purpose, we used the domains of the computerized multiphasic interactive neurocognitive system (CMINDS) scores including speed of processing, attention/vigilance, working memory, verbal learning, visual learning, and reasoning/problem solving. Further details of CMINDS and preprocessing steps can be found in (van Erp et al., 2015).

### 2.8. Statistical analysis: dynamic alterations among patients with schizophrenia

The spatiotemporal variations of brain networks were evaluated by comparing group differences between patients with SZ and healthy subjects. For each statistical comparison, we used a general linear model (GLM) that included age, gender, site, and a mean framewise displacement (meanFD) as covariates. meanFD was added as a covariate to mitigate against the effects of motion (Power et al., 2012). To evaluate the relationship between energy indices and CMINDS scores, the correlation analyses were conducted separately for healthy subjects and patients with SZ after regressing out age, gender, site, and meanFD. CMINDS scores and energy indices were mean-centered for each group to mitigate group differences in either CMINDS scores and/or energy indices introduced into the correlations. The group difference in correlation coefficient of the relationship between energy indices and CMINDS scores was also evaluated between the two groups. Outliers were removed using the scaled median absolute deviation, and *p*-values of all statistical analyses were corrected for multiple comparisons using the 5% false discovery rate (FDR) (Benjamini and Hochberg, 1995).

## 3. Results

### 3.1. Large-scale brain networks

We applied spatial ICA with 20 components on rsfMRI data from 309 individuals. The ICA algorithm ran several times and clustering the resulting components revealed a high level of compactness (a cluster quality index greater than 0.8), indicating the reliability of the independent components (Himberg et al., 2004). Twelve independent components were identified as neuronal activity related components, or brain networks (Erhardt et al., 2011a), based on their temporal and spatial properties and knowledge from previous studies. The time courses of the selected components are dominated by low-frequency fluctuations, which were evaluated using the dynamic range and the ratio of low-frequency to high-frequency power (Allen et al., 2011). Their spatial maps have significant overlap with gray matter, their peak of activations fall within gray matter, and display low spatial overlap with known ventricular, motion, and susceptibility artifacts components (Allen et al., 2011). Furthermore, the selected components should have high spatial similarity with one of the established intrinsic connectivity networks (Allen et al., 2011; Beckmann et al., 2005; Damoiseaux et al., 2006; Fox et al., 2006; Iraji et al., 2016; Smith et al., 2009; Yeo et al., 2011; Zuo et al., 2010). The identified brain networks are the auditory, cerebellar, default mode, (dorsal) attention, left and right frontoparietal, somatomotor, language, salience, subcortical, primary visual, and secondary visual networks (Figure 2). After identifying the brain networks, the time courses of individual networks were used to calculate their dCMs (one dCM per window for each brain network).

**Figure 2.**
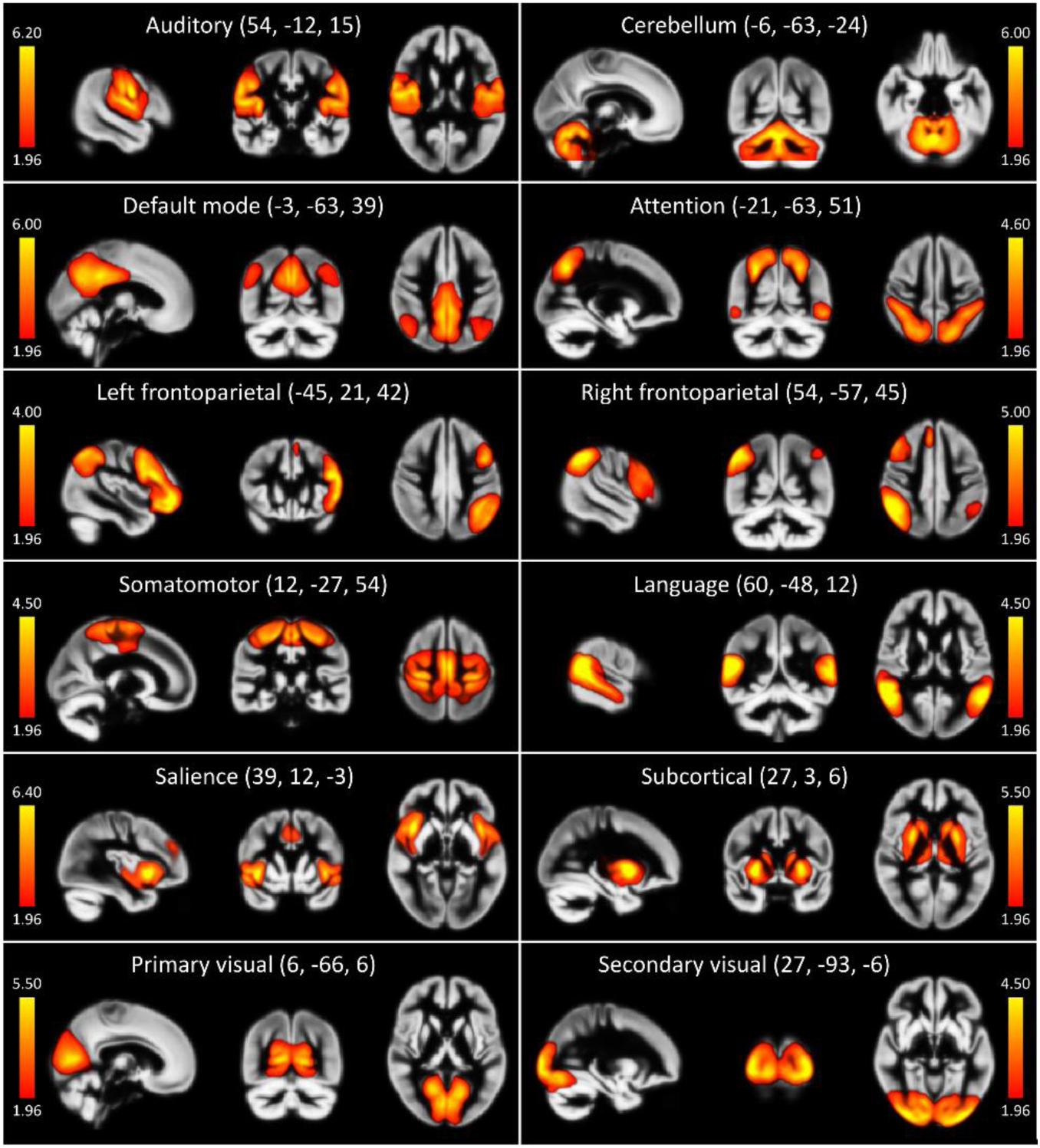
Static group spatial functional connectivity maps of the twelve brain networks obtained from spatial independent components analysis. Spatial maps are plotted as z-score and thresholded at Z > 1.96.

### 3.2. Spatial state evaluation

To evaluate the time-varying information of brain networks, k-means clustering was applied on the dCMs of each network to group them into a set of distinct spatial patterns called spatial states. This allows us to visualize and quantify the variations in spatial patterns of brain networks. An example of variations in spatial patterns of brain networks and the regions associated with them can be seen in Figure 3, in which three selected spatial states for each brain network represented as red, blue, and green additive color overlays. Regions in white indicate association to a given network in all three states. For instance, the posterior cingulate cortex (PCC) is the central hub of the default mode (Andrews-Hanna, 2012; Leech and Sharp, 2014) and expected to be part of the default mode all of the time, which is confirmed by being involved in all three states as well as the fourth state, which is not included in Figure 3. However, the thalamus is associated with the default mode in one of the three states (represented in red), and the frontal regions are involved in either one (represented in blue or red) or two states (represented in purple). These findings could explain inconsistencies in previous findings regarding the membership of certain regions to brain networks. For instance, different patterns for the default mode have been observed (Andrews-Hanna et al., 2010; Braga and Buckner, 2017; Fox et al., 2005). While most of the original studies did not report the thalamus as part of the default mode (Buckner et al., 2008; Fox et al., 2005; Greicius et al., 2003), recent studies have shown conflicting results (Lee and Xue, 2018; Shirer et al., 2012; Wang et al., 2014). Moreover, different parts of the frontal and prefrontal lobes have been reported as part of the default mode across studies (Braga and Buckner, 2017; Buckner et al., 2008; Damoiseaux et al., 2006; Garrity et al., 2007; Shirer et al., 2012). Similarly, different patterns of regions memberships have been observed across other brain networks (Braga and Buckner, 2017; Damoiseaux et al., 2006; Zuo et al., 2010). Therefore, we suggest that different regions are associated with brain networks at different time points, and only overall patterns of brain networks during data acquisition are identified in the static analysis.

**Figure 3.**
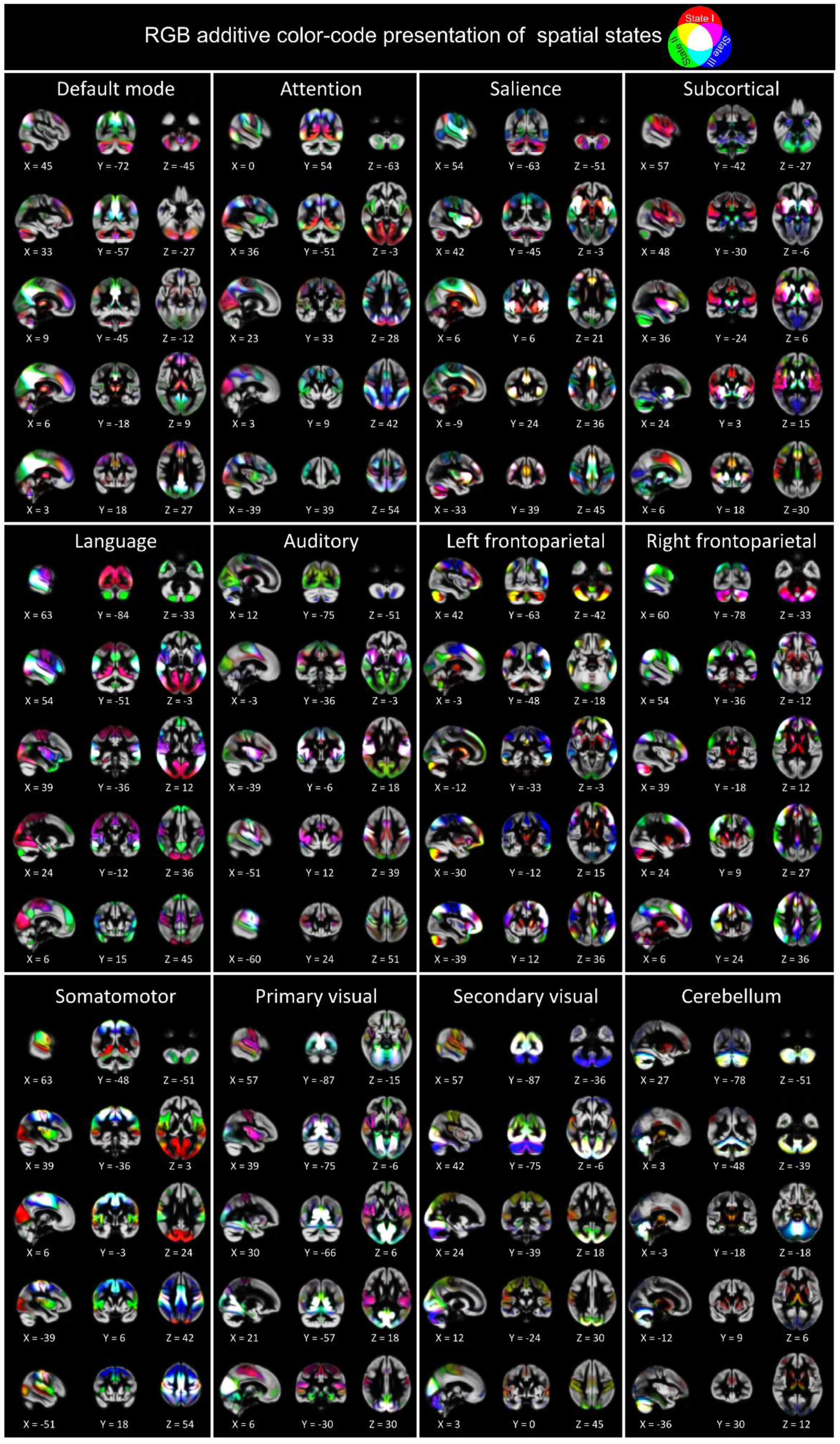
RGB additive color-code presentation of three arbitrarily-selected spatial states for brain networks. Red, blue, and green represent the strength of three spatial states. Thus, for instance, white represents the areas in which the brain network is strong in all three spatial states, and yellow shows strong association in Red and Green spatial states. It is worth mentioning that the spatial states of different brain networks were identified independently; therefore, for example, the first spatial state of networki is not correspondent with the first spatial state of networkj (i ≠ j).

**Anticorrelated brain networks**: Evaluating the spatial chronnectome using the dCMs also provides new information relating to anticorrelated brain networks, i.e., negative associations between brain networks. Previous static studies have observed negative associations between brain networks and their associated regions (Allen et al., 2011; Fox et al., 2005; Fox et al., 2009; Uddin et al., 2009). For instance, regions of different brain networks (including salience and sensorimotor) are suggested to be negatively correlated with the default mode regions (Allen et al., 2011; Fox et al., 2005; Fox et al., 2009; Uddin et al., 2009). Our analysis reveals that different, anticorrelated patterns occur at different momentsin time. For example, each of previously reported networks and its associated regions become negatively associated with the default mode for a specific spatial state, and there are moments in which no negative associations exist (Figure 4). In other words, anticorrelative relationships identified across previous default mode static analyses all exist, but in differing segments of time. We further observed new anticorrelated patterns across different networks including in left and right frontoparietal, salience, somatomotor, and secondary visual (Figure S2). This finding emphasizes the importance of time-varying properties that may not be fully captured during static analysis.

**Figure 4.**
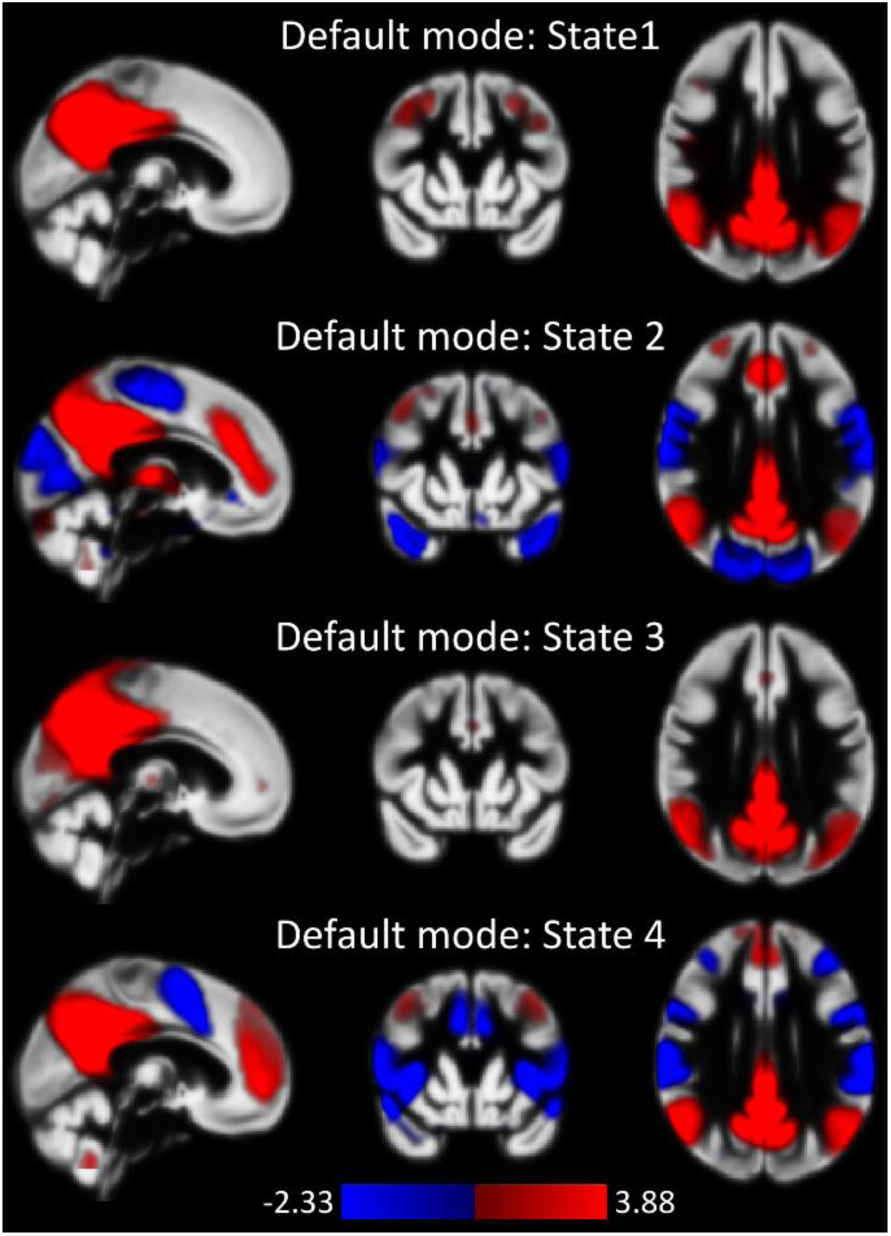
The spatial states of the default mode. Hot and cold colors represent positive and negative associations to the default mode, respectively. The results show that sensorimotor areas are anti-correlated with the default mode during State 2, and the salience network is anti-correlated with the default mode during State 4. Importantly, States 1 and 3 do not exhibit an anticorrelative relationship with the default mode.

**Cerebellar contribution:** Despite the important role of the cerebellum, it is often overlooked in brain network analysis. One reason is that the cerebellum is not usually recognized as an integral part of the connectome, with some exceptions, across studies using static analysis. Our analysis reveals significant contributions of the cerebellum to multiple brain networks, but these contributions are not constant over time (see Figure 3). Different patterns of cerebellar contribution emerge at particular timepoints or states. This highlights a challenge in detecting the role of the cerebellum in brain networks in static analysis. Overall, two major patterns are 1) primarily negative associations between cerebellar regions with sensorimotor networks (e.g. including somatomotor, auditory, and visual networks), and 2) positive associations of cerebellar regions with the subcortical and left and right frontoparietal networks.

**Brain networks are not isolated:** Studying spatial chronnectomes through spatial states reinforces our understanding that brain networks are not isolated, and there is strong cross talk between “isolated” brain networks. Regions assigned to one network using static analysis are also involved with other networks at particular points in time. This is observed across all networks but is more dominant in sensorimotor networks including visual, somatomotor, and auditory networks (Figure 3). This finding further suggests that brain networks sometimes merge, or inter-network coupling increases, consistent with a dynamic interplay between segregation and integration. Therefore, evaluating the interaction between networks and their associated regions using this approach can shed new light on the multifactorial role of large-scale brain networks.

**Statistical comparison:** The dCMs of brain networks were compared between healthy subjects and patients with SZ. We initially hypothesized that networks’ dCMs would allow us to detect nuanced alterations in brain networks in patients with SZ relative to healthy subjects which would be lost and remain undetected during static analyses. This hypothesis was evaluated first by comparing the results of voxel-wise comparisons using spatial states (obtained by applying K-means clustering on the dCMs of each brain network) and static connectivity (spatial map obtain by applying sICA on the data of all time points) analyses. The results of voxel-wise comparisons are presented in Figure 5(A). Both static and spatial states approaches reveal group differences in the cerebellar, subcortical, language, and salience networks. The pattern of differences in these networks was similar between static and spatial states analysis. Static comparison revealed lower static functional connectivity in patients with SZ compared to healthy subjects across these networks except for the putamen in the subcortical network. Similarly, spatial states analyses detected decreases in dynamic couplings across the same networks with the same exception in the subcortical network. In contrast, different and larger regions within networks were found to be altered in patients with SZ compared to healthy subjects in the spatial state analysis compared to the static analysis. Additionally, spatial state analysis shows similar patterns of differences between patients with SZ and healthy subjects in the auditory, primary and secondary visual, somatomotor, default mode, (dorsal) attention, and left frontoparietal networks (Figure 5(A)). The similarity between static and state analysis patterns further supports the idea that alterations observed in state analysis driven by group differences between patients and healthy subjects. This suggests state analyses detect nuanced alterations within patient groups absent in static analyses. For instance, an alteration in a small region of the primary visual network can also be seen in static analyses if we apply less restrictive criterion i.e., using smaller cluster size threshold (Figure S3), which reiterates the loss of important nuanced information through static analysis. Previous studies suggest these networks and associated areas are affected in patients with SZ (Baker et al., 2014; Calhoun et al., 2009; Garrity et al., 2007; Jafri et al., 2008; Zeng et al., 2018), which indicates the ability of the approach to both detect known patterns of alterations and also identify novel patterns of patient-based alterations.

It is important to highlight some of the regions within a network that show differences in the state analysis were not recognized as being part of the same network in a static analysis. In other words, a region that is part of Network *i* at State *k* and assigned to the Network *j* in the static analysis display significant differences between healthy subjects and patients with SZ. Figure 5(B) represents a summary of spatial state voxel-wise comparisons and their static labeling. The Y-axis includes the spatial states of the networks illustrating significant differences between healthy subjects and patients with SZ. The X-axis indicates the static network assignment of the regions that show significant differences. For instance, the lingual gyrus is assigned to primary visual network in static analysis, but it is also part of (associated with) at State 1 of the language domains. This region shows a reduction in its association with State 1 of the language domain in patients with SZ. This emphasizes the importance of capturing dynamic information about network integration.

**Figure 5.**
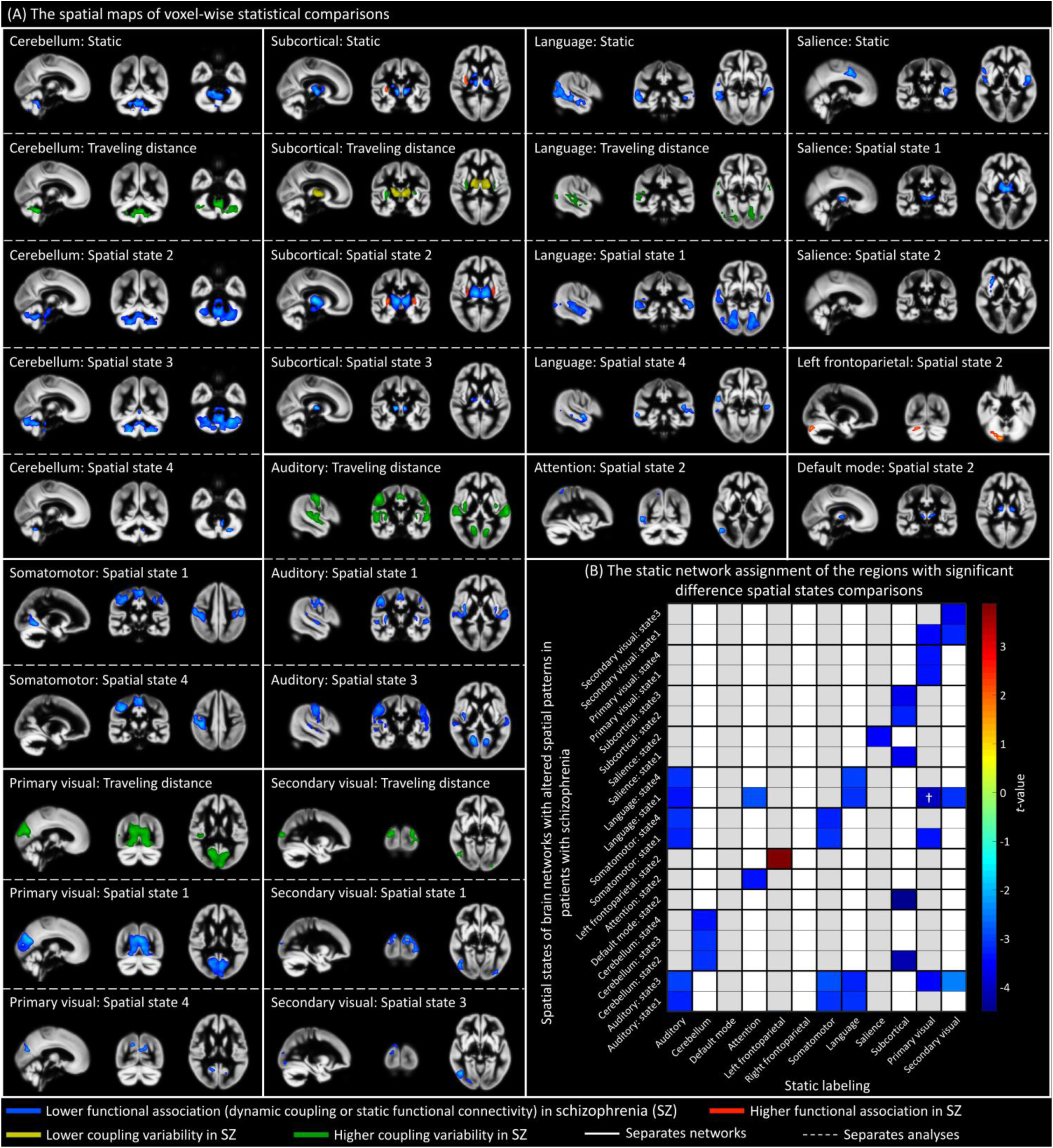
Voxel-wise statistical comparisons between healthy subjects and patients with schizophrenia (SZ). Only comparisons that show significant differences after multiple comparison corrections are presented here. (A) Spatial maps displaying significant differences between healthy subjects and patients with SZ. Blue and red colors represent lower and higher associations of regions to the networks in patients with SZ relative to healthy subjects, respectively. Yellow and green colors indicate lower and higher coupling variability, i.e., variation over time measured by the L1 norm distance, in patients relative to healthy subjects. Networks are separated with solid white colors, while different types of analyses including static functional connectivity, coupling variability (L1 norm distance), and spatial states are separated by dashed white color lines. (B) Summary of voxel-wise comparisons of spatial states. The Y-axis includes the list of the spatial states of the networks with significant differences between the two groups. The *t*-values for differences between patients with SZ vs. healthy subjects. Cold and hot colors indicated transient reduced and increased association (network coupling) in patients with SZ relative to healthy subjects, respectively. The X-axis indicates the static network labeling of the regions that show significant differences in state analysis. For instance, the cell labelled with “†” represents the voxels that show a significant difference with reduced dynamic network coupling in patients with SZ (*t*-value) in State 1 of the language network (Y-axis). These voxels are assigned to (labeled as) the primary visual network in the static analysis (X-axis).

While the spatial states of brain networks provide important details of their spatial dynamic, their temporal profiles provide further information regarding the temporal dynamic nature of brain networks. Statistical analyses of mean dwell time and fraction time showed statistically significant differences between patients with SZ and healthy subjects across all networks (Figure 6). In patients with SZ, networks tend to spend more time in spatial states which have high correlations, negative or positive, with sensorimotor regions, particularly within primary visual areas. They include State 1 of the right frontoparietal network, State 3 of the primary visual network, State 1 of the somatomotor network, State 2 of the subcortical network, State 2 of the default mode network, State 2 of the cerebellar network, State 1 of the salience network, State 1 of the left frontoparietal network, State 2 of the attention network, and State 1 of the language network. We also observed certain networks spent more time in the spatial states with high (negative or positive) correlations with the default mode regions in healthy subjects compared to patients with SZ.

**Figure 6.**
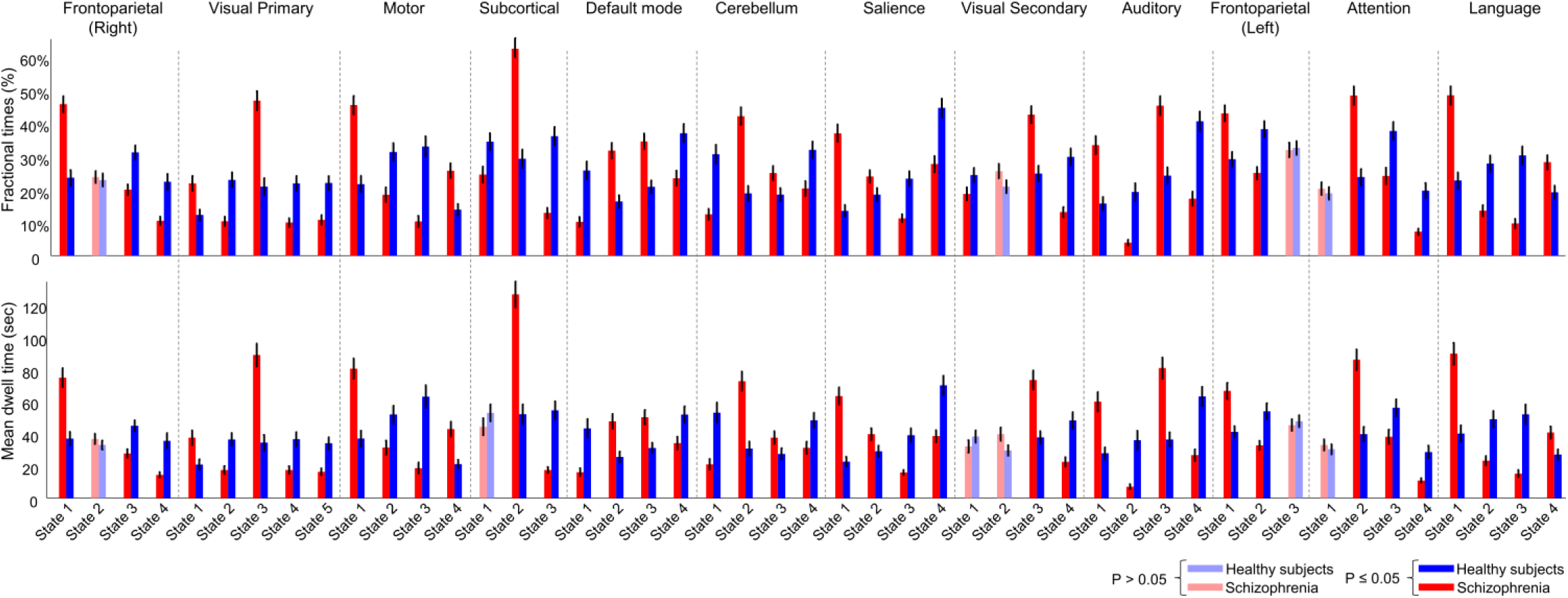
The ability of dynamic temporal indices calculated from spatial states to distinguish different cohorts. Two indices including fractional time and mean dwell time were calculated for spatial states of brain networks and compared between healthy subjects and patients with SZ. Statistically significant differences after multiple comparison corrections were observed between patients with SZ and healthy subjects. The results suggest the ability of the approach to detect patient-based alterations.

### 3.3. Spatial variation evaluation

#### Coupling variability map

Brain networks are spatially fluid, and this can be captured and evaluated via calculating coupling variability maps over time. Figure 7 shows variations in the dynamic coupling of associated voxels to a given network over time. Green represents coupling variability, indicating variations in dynamic coupling of each voxel to a given network over time. Red represents static functional connectivity strength. For instance, our results show the PCC that is always associated with the default mode has lower variation over time. However, the thalamus and frontal areas reveal higher variations over time. Evaluating variation in regions association to brain networks can provide further information about their roles in brain networks. Performing voxel-wise comparison of coupling variability maps reveal significant differences between healthy subjects and patients with SZ in cerebellar, subcortical, language, auditory, and primary and secondary visual networks (Figure 5(A)). Thus, we observed higher coupling variability, in addition to lower network coupling strength, among patients with SZ. Collectively, these results suggest both coupling strength and variability are altered in patients with SZ.

**Figure 7.**
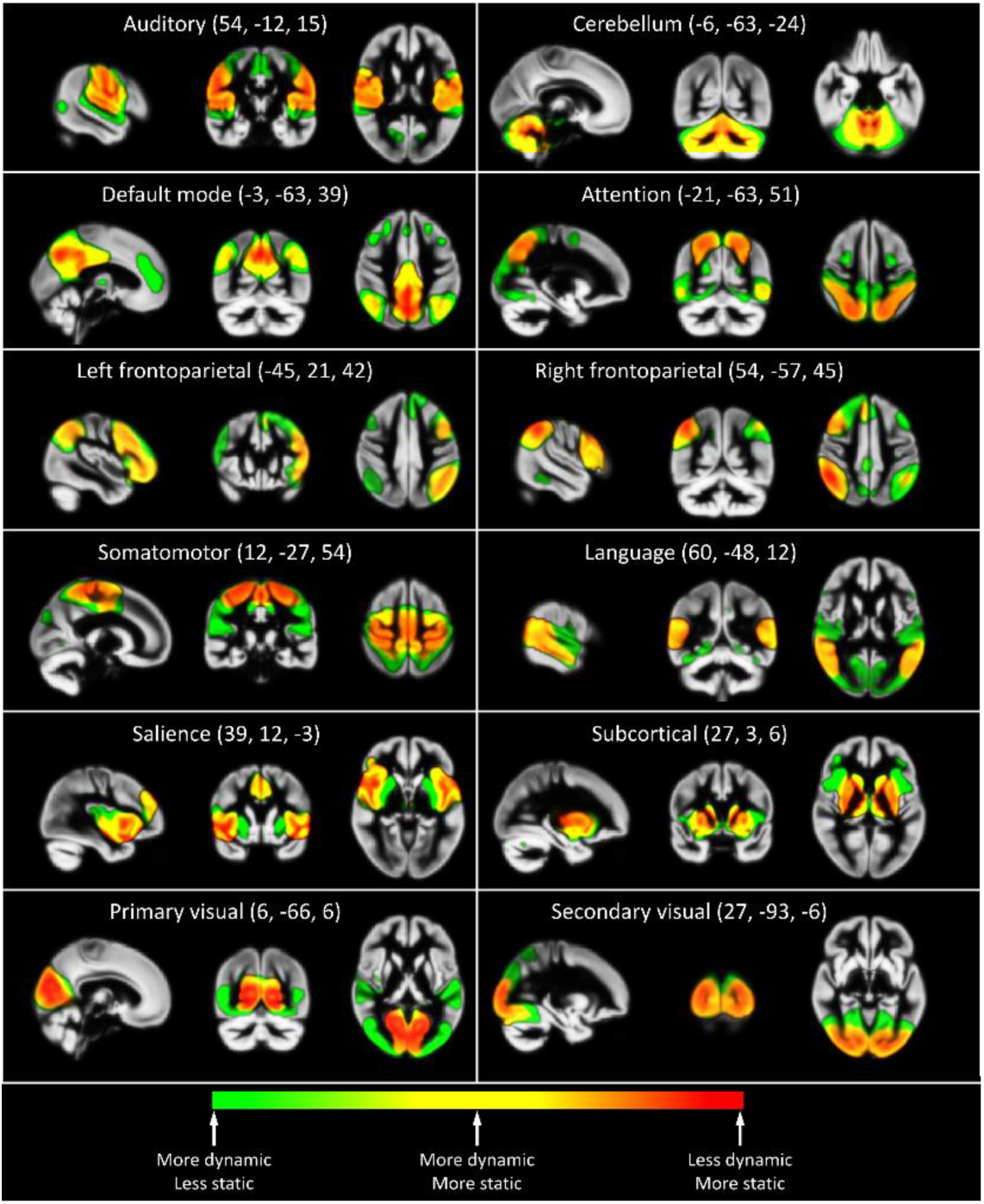
Additive color-code representation of networks’ coupling variability. Green represents coupling variability estimated by the L1 norm distance of variations in the membership (pairwise correlation) of each voxel to a given network over time. Red represents static functional connectivity strength. Thus, Yellow represents the regions with both high coupling variability over time and static strength. The figure indicates extensive variations in spatial patterns of brain networks. Previous work typically ignores levels of variability and the degree to which a voxel contributes to a given network over time, which is captured using this method.

#### Spatiotemporal transition matrix

While voxel-wise analysis is important to assess the spatial variations of brain networks, the spatiotemporal transition matrix can quantify and summarize dynamic spatiotemporal properties of individual brain networks, which are essential to understanding the dynamic characteristics of brain networks. Because of the smoothing effect of the sliding window, the spatiotemporal dynamic properties become easier to distinguish as the interval value increases. However, the pattern of transitions is consistent across interval values (See examples in Figure S4). In Figure 8, we present the examples of findings for the interval value of 30 (the size of the interval = 30×TR), which is the transition between two windows with almost no overlap (Figure 8(A)). Evaluating the spatiotemporal dynamic properties allows us to detect changes in the brain networks such as the default mode which were imperceptible in our static analysis. Performing statistical comparisons on the elements of the spatiotemporal transition matrix display its ability to differentiate healthy subjects and patients with SZ (Figure 8(B)). Patients with SZ have higher transitions in higher bins, i.e. bins with higher dynamic coupling value, and healthy subjects have higher transitions in lower bins, with the exception of the default mode, which shows the opposite pattern. Significant differences were observed between healthy subjects and patients with SZ in all networks expect the left and right frontoparietal networks. Examples of the comparisons for all possible interval values for multiple networks are included in Figure S5, which display similar patterns for different interval values. The spatiotemporal transition matrix can further be utilized to extract abstract global measures to summarize the dynamic properties of each network. For instance, evaluating spatiotemporal uniformity using the energy index shows a significant difference between the two groups in which healthy subjects demonstrated higher spatiotemporal uniformity (i.e., lower energy index) (Figure 8(C)).

**Figure 8.**
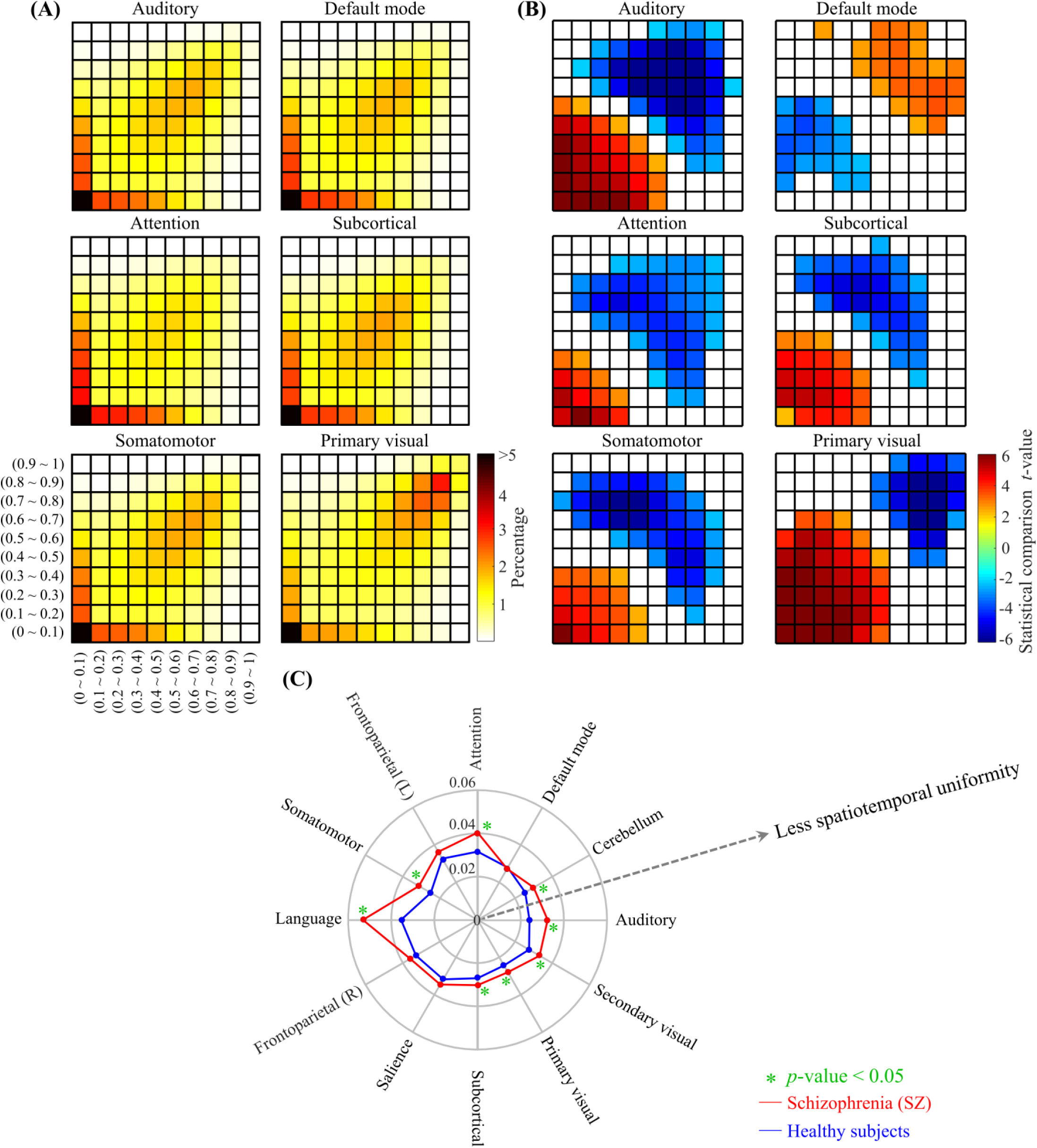
Examples of spatiotemporal transition matrices and statistical analysis. (A) The average spatiotemporal transition across all subjects. The spatiotemporal transition matrix for each network summarizes the variation of the network’s dCMs by discretizing the dynamic coupling values into ten bins. The value in each array indicates the percentage of the transition compared to the total number of transitions using warm colors. For instance, if the number of transitions were uniform, the value of each array would be 1% because there are 100 arrays in the transition matrix. (B) *t*-statistics for group comparisons by diagnosis. Blue (cold) and red (hot) colors represent lower and higher transition values in patients with SZ compared to healthy subjects, respectively. (C) Energy index comparison, with greater spatiotemporal uniformity towards the center of the chart. The energy index was measured for the spatiotemporal transition matrix and compared between healthy subjects and patients with SZ. Blue and red colors represent healthy subjects and patients with SZ, respectively. Green asterisks indicate the statistical significant differences between the two groups.

Evaluating the relationship between CMINDS and the energy indices display significant correlations between the energy index of the subcortical domain from the imaging data and attention/vigilance CMINDS domain in the healthy subjects (ρ = -0.27; *p* < 0.0011, FDR = 0.049) but not patients with SZ (ρ = 0.12; *p* < 0.1561, FDR = 0.962). Furthermore, the group difference of the correlation between the energy index of the subcortical domain from the imaging data and Attention/Vigilance CMINDS domain approached statistical significance (*p* < 0.0012, FDR = 0.059).

## Discussion

Static analysis of fMRI data (i.e. computing correlations based on all timepoints) has provided important information about the brain; however, the assumption that resting brain activity can be represented by static activity across time is a gross oversimplification which may obscure the true dynamic nature of the brain. Motivated by the recent discovery of the ability of fMRI to capture dynamic, time-varying information of brain connectivity, we propose a spatial chronnectome approach which examines the variations in the spatiotemporal coupling of networks at the voxel level. The findings of our study offer unprecedented evidence for the spatially fluid behavior of inter- and intra-network brain relationships and emphasize the dynamic interplay between information segregation and integration across the brain. Our approach affirms that brain networks evolve spatially over time by capturing spatiotemporal variations within brain networks. For instance, our approach identified variability in the brain network membership of a given brain region over time.

For discussion purposes, regions associated with a given brain network can be divided into two categories. The first includes regions which are repeatedly or occasionally reported to be parts of a given network in static analysis. The second category contains brain regions known to be parts of other networks in static studies based on previous research. Findings related to the first category explain the inconsistent observations regarding the brain network spatial patterns. These findings indicate that the spatial chronnectome, i.e., the temporal variations of the coupling patterns of brain networks, is the reason for the observed inconsistency in the spatial patterns of brain networks and the variability in brain regions’ memberships. For example, thalamus and frontal regions show high variations in their associations to the default mode, even dissociating from the default mode at particular times. The PCC, however, showed a more robust and constant association to the default mode over time, which may be related to its role as the central hub (core) of the default mode. The small amount of variation/variability of the PCC association with the default mode suggests that the cores of brain networks have smaller variations in their dynamic couplings to the associated networks over time. Categorizing regions based on their time-varying associations with brain networks, and evaluating the multifactorial roles and relationship between them can provide new information about the interaction within and between network.

It is worth mentioning that there is both a high-level of similarity and major differences between the spatial states and the interdigitated networks observed in previous single-subject analysis (Braga and Buckner, 2017) which can be seen, in our opinion, as an alternative interpretation for the observed inconsistency in the spatial patterns of brain networks. The concept of the spatial chronnectome within brain networks does not exclude the existence of a set of parallel networks within each large-scale network, as these also can be identified via brain dynamic analysis. Because a brain region’s association to a given large-scale network varies over the time, the associated parallel networks can be captured using the time points during which they contribute in the dominant patterns within the given large-scale networks.

The second category highlights that brain networks are not isolated and exhibit significant cross talk. Regions in different networks join other networks and dissociate from them over time. This pattern of integration and segregation occurs across various regions and all investigated networks. This finding helps to extend the classical view of information processing in the brain and detect what we are currently missing about the functions of brain regions. An example of this claim is our observation that the primary visual area transiently associated with multiple networks which do not typically include visual areas. In our analysis, the primary visual area demonstrates significant positive or negative associations to at least one state of all identified brain networks suggesting its major role in network cross talk and various brain functions. This finding challenges the classic view of primary visual area role as responsible for receiving and delivering visual information from retinal input. However, there is significant evidence to support the role of the primary visual area in other brain functions. While the primary visual area receives most of the retinal input (90%) (Sincich et al., 2004), neuronal tracing and neuronal recording investigations demonstrated feedback connections between primary visual and many cortical areas (Bullier et al., 2001; Felleman and Van Essen, 1991; Hupe et al., 2001). For instance, the primary visual area receives information from a wide range of sensory and non-sensory cortices such as the primary auditory, parietal and frontal cortices (Markov et al., 2011). Moreover, transcranial magnetic stimulation (TMS) interference on the primary visual area introduced impairments to working memory processing (van de Ven et al., 2012). However, the classic view of the primary visual cortex can be well studied by fMRI because fMRI can detect changes associated with higher cognitive function and indirect functional connectivity. fMRI research provides the striking evidence that the primary visual cortex is involved in higher cognitive functions (Bressler et al., 2013; Harrison and Tong, 2009; Lars, 2010; Muckli and Petro, 2013; Roelfsema and de Lange, 2016). To conclude, variability in a given region’s association to a given brain network highlights a dynamic interplay between segregation and integration, providing a new perspective on the function of well-known brain areas. Our results suggest evaluating time-varying properties of brain network interplay is thus a vital issue for future research to understand the multifactorial role of brain networks.

Studying patterns of brain networks using spatial states also provides striking findings regarding anticorrelative patterns of brain networks. Similar to our finding that different brain regions become involved with different brain networks over time, various anticorrelative relationships were detected at different time points. Using a spatial state analysis, the default mode shows anticorrelated relationships with sensorimotor and salience networks at States 2 and 4 respectively, but the anticorrelated pattern attenuates in States 1 and 3. Our findings suggest anticorrelative relationships are not limited to the default mode. We identified a new set of anticorrelated patterns for various networks and illustrate, for the first time, anticorrelative relationships occur at specific moments rather than persisting over time. This is an intriguing finding which suggests the need for new investigations and potential revision to proposed causality or modulatory relationships between brain networks such as between the salience and default mode.

Studying dCMs of brain networks also accentuates the role of the cerebellum in different networks. The cerebellum has widespread polysynaptic connections to the cerebral cortex which all pass through the thalamus (Buckner et al., 2011; Kelly and Strick, 2003; Krienen and Buckner, 2009; Strick et al., 2009). This phenomenon can explain the role of the cerebellum in a wide range of motor and cognitive functions (Lars, 2010; Muckli and Petro, 2013; Stoodley and Schmahmann, 2009). While the polysynaptic connections make reconstructing the topography of cerebro-cerebellar connections by anatomical methods relatively difficult, the ability to measure indirect connectivity (functional connectivity) via fMRI makes it an effective way to map cerebro-cerebellar connections (Krienen and Buckner, 2009; O’Reilly et al., 2010). Our observations, similar to previous fMRI studies, demonstrate stronger contralateral connectivity patterns between cerebellum and cerebral cortex compared to ipsilateral connectivity patterns, which is consistent with known contralateral polysynaptic connections between cerebral cortex and the cerebellum (Buckner et al., 2011; Kelly and Strick, 2003; Krienen and Buckner, 2009; O’Reilly et al., 2010). In general, previous rsfMRI studies agree on the relationship of the cerebellum with the thalamus, motor area, and regions involved with the frontoparietal networks (Krienen and Buckner, 2009; O’Reilly et al., 2010). Our dynamic network analysis corroborates these results.

Despite the unique ability of rsfMRI to measure cerebro-cerebellar connectivity, the role of the cerebellum in brain network analysis is often overlooked. This is in part because it is not usually recognized as an integral part of the connectome, and its static functional connectivity to brain networks is typically weaker compared to cortical connectivity limiting the visibility of the cerebellum due to limitations in statistical power. In earlier work, Buckner and his colleagues estimated specific cerebellar topography patterns for cerebral networks by assigning each cerebellar voxel to a network with the highest correlation values (Buckner et al., 2011). Although their approach had limitations preventing concrete conclusions, their work provides a striking result regarding different functional connectivity patterns across the cerebellum. Using a dynamic analysis, we observed intriguing findings suggesting the evaluation of time-varying contributions of cerebellar regions to different brain networks conjointly provide a great deal of knowledge about the role of the cerebellum in functional networks. Our analysis showed positive associations between cerebellar regions and the subcortical network, which may be explained by the relay role of the thalamus in cerebro-cerebellar connections. We also observed positive associations between cerebellar regions and both the left and right frontoparietal networks that relate to fronto-cerebellar circuitry (Kelly and Strick, 2003; Krienen and Buckner, 2009; O’Reilly et al., 2010). For the first time, we observe negative associations between cerebellar regions and certain sensorimotor networks. These include somatomotor, auditory, and visual networks. This negative association could be related to modulating the connection between the thalamus and cortex in cerebellar thalamic cortical circuits.

It is also worth mentioning that functional relationship between the cerebellum and somatosensory and motor/premotor cortex has been frequently reported, but there is some disagreement over the relationship between the cerebellum and both the primary auditory and visual cortices (Krienen and Buckner, 2009; O’Reilly et al., 2010). O’Reilly et al. (O’Reilly et al., 2010) suggested the relationship between the cerebellum and primary auditory and visual areas probably reflects the importance of visual and auditory information in motor control which was demonstrated in earlier experiments. Interestingly, the motor area is positively associated with the states of the primary visual and auditory networks that have a negative association with the cerebellum. This also emphasizes the importance of our finding that different networks have cross talk.

Our approach emphasizes distinct features in both the spatial and temporal realms. Regarding spatial patterns of brain networks, the spatial chronnectome analysis allows us to detect atypical brain patterns in patients with SZ that cannot be detected using a static analysis including statistical alteration in dCMs. Moreover, spatial chronnectome analysis detects important variations in temporally fluid patterns. For example, the influence of dynamic alterations in the static analysis can be small, and therefore changes networks coupling would not be detected in static analysis due to a reduction in the statistical power. With respect to temporal dynamic properties, preliminary investigations reveal significant differences in both mean dwell time and fraction time between patients with SZ and healthy subjects. Additional independent studies will be needed to comprehensively investigate the temporal properties of brain networks through networks’ state and meta-state indices. Furthermore, our approach provides a unique opportunity to investigate the variations of networks’ couplings which is not feasible in static analysis. In our analysis, we observed overall higher coupling variability and lower network coupling strengths among patients with SZ, which is aligned with the dysconnectivity or disconnection hypothesis of schizophrenia (Friston, 1998). Reduced functional connectivity has been constantly reported in patients with SZ (Damaraju et al., 2014; Erdeniz et al., 2017; Gavrilescu et al., 2010; Skudlarski et al., 2010; Vercammen et al., 2010), and higher fluctuations of brain connectivity within brain networks in SZ could be related to the brain’s effort to compensate for dysconnectivity and/or unbalancing of brain circuitries (Cazorla et al., 2015).

While the proposed approach can capture the spatial fluidity of networks’ couplings, new indices are also needed to quantify the spatiotemporal of individual brain networks. For this purpose, the spatiotemporal transition matrix and associated features were introduced to summarize the time-varying properties. Statistical comparisons between healthy subjects and patients with SZ reveals statistical significant differences which were consistent across interval values. With the exception of the default mode, the patterns of alterations are similar across networks in which patients with SZ have higher transitions in higher dynamic coupling values, while healthy subjects display higher transitions in lower dynamic coupling values. The opposite pattern of the default mode could be related to its activity pattern in relation to other networks and mental and physical activities. It is known that static functional connectivity within the default mode decreases as the static functional connectivity of the other networks increases. For instance, during a task, the connectivity value within the default mode reduces. Thus, it is expected that variations in lower dynamic coupling values of the default mode are associated with variations in higher dynamic coupling of other networks. Significant differences between healthy subjects and patients with SZ were also observed in networks’ spatiotemporal uniformity. Future work should also examine how spatiotemporal dynamic information can improve classification accuracy of patients into diagnosis categories. More importantly, significant associations were detected between spatiotemporal uniformity indices obtained from the spatiotemporal transition matrix and attention/vigilance cognitive domains of CMINDS. Previously, van Erp et. al., found patients with SZ revealed significantly large impairments in the speed of processing and the attention/vigilance of CMINDS compared to healthy subjects on the, more so than other CMINDS domains (van Erp et al., 2015). As such, our findings of strong difference in the links between subcortical and CMINDS attention/vigilance domain may reflect a true disruption in subcortical domains relative to attention/vigilance in patients with SZ, but lack the statistical power for confirmation. Future work is needed to identify the nature of the relationship between spatiotemporal indices and neuropsychological variables in both SZ and other patient groups, but our preliminary results demonstrate promising relationships between the spatial chronnectome metrics and cognition.

**Limitations and future directions:** The proposed approach is utilizing the sliding window (Allen et al., 2014; Damaraju et al., 2014; Sakoglu et al., 2010), which is the most commonly used approach to study time-varying properties of brain networks. Previous analysis suggested that data length of 30-60 seconds is a good choice to successfully capture the dynamic properties (Allen et al., 2014; Preti et al., 2017) as well as to estimate cognitive states (Shirer et al., 2012). While we chose a window size of 60 seconds to follow previous recommendations, we highlight the importance of capturing the dynamic information of BOLD signals to its full potential, i.e. up to the maximum frequency that exists in the data (Trapp et al., 2018; Vidaurre et al., 2017; Yaesoubi et al., 2018). Therefore, the approach should be improved to capture the full amount of time-varying information in the data. The other drawback of this work is the spatial resolution of the data, which limits the dynamic spatial specificity. Spatial resolution and smoothing induce blurring, which can be detrimental for capturing time-varying properties of brain networks. This effect has a more severe impact when brain regions with very distinct functional roles, like sub-regions of the cerebellum, are located in close proximity to one another within a small area. Data with higher spatial and temporal resolutions can provide better insight into spatiotemporal variations of brain functional organizations. Moreover, using surface-based registrations for high spatial resolution data instead of volume-based registration could potentially enhance functional specialization.

In this study, we investigated the spatiotemporal variations of the brain networks, i.e. spatial independent functional organizations; however, another set of functional organizations can be obtained by assuming temporal dependency (Calhoun et al., 2001b; Smith et al., 2012). Temporal independence may be better suited for the proposed approach, as it does not assume spatial stationarity in the first step of the analysis. In other words, using spatially independent networks carries the same contradictory assumption regarding the spatial maps that exists in spatio-temporal (dual) regression analysis (Erhardt et al., 2011b). Thus, future studies with higher temporal resolution should be used to investigate the spatial chronnectome of temporally independent networks. Finally, the proposed approach is the first step toward enhancing our understanding of spatiotemporal variations of brain functional organizations at the voxel level. Further work needs to be conducted to transform the approach to an established framework to investigate the spatial chronnectome of functional organizations across different cohorts (e.g. patients and healthy subjects).

## Conclusion

The proposed approach provides a new framework to study the spatial chronnectome of brain functional organizations. Despite the limitations of our analysis/acquisition approach, such as spatial and temporal resolutions and sliding window restrictions, the findings of this study elucidate the spatiotemporal variations of brain networks. Major findings of the study include: 1) highlighting spatially fluid behavior of intra- and inter-network relationships, underlying a dynamic interplay between segregation and integration of information; 2) explaining broad-spectrum of inconsistencies in findings of static analysis; and 3) extracting detailed information and nuanced alterations of brain networks which gets lost and remains undetected using static analysis. Furthermore, new indices are introduced to evaluate spatiotemporal variations in brain functional organizations such as brain networks. Preliminary assessments of the approach using healthy subjects and patients with SZ demonstrate the ability of the approach to obtain new information of brain function and detect alterations among patients with SZ. Further investigation should be performed to evaluate the ability of the approach to study spatiotemporal variations of brain functional organizations.

## Acknowledgments

This work was supported by grants from the National Institutes of Health grant numbers 2R01EB005846, R01REB020407, and P20GM103472; and National Science Foundation (NSF) grant 1539067 to Dr. Vince Calhoun. We thank Srinivas Rachakonda and Eswar Damaraju for their input.

## References

Allen, E.A., Damaraju, E., Plis, S.M., Erhardt, E.B., Eichele, T., Calhoun, V.D., 2014. Tracking whole-brain connectivity dynamics in the resting state. Cereb Cortex 24, 663–676.

Allen, E.A., Erhardt, E.B., Damaraju, E., Gruner, W., Segall, J.M., Silva, R.F., Havlicek, M., Rachakonda, S., Fries, J., Kalyanam, R., Michael, A.M., Caprihan, A., Turner, J.A., Eichele, T., Adelsheim, S., Bryan, A.D., Bustillo, J., Clark, V.P., Feldstein Ewing, S.W., Filbey, F., Ford, C.C., Hutchison, K., Jung, R.E., Kiehl, K.A., Kodituwakku, P., Komesu, Y.M., Mayer, A.R., Pearlson, G.D., Phillips, J.P., Sadek, J.R., Stevens, M., Teuscher, U., Thoma, R.J., Calhoun, V.D., 2011. A baseline for the multivariate comparison of resting-state networks. Front Syst Neurosci 5, 2.

Andrews-Hanna, J.R., 2012. The brain’s default network and its adaptive role in internal mentation. Neuroscientist 18, 251–270.

Andrews-Hanna, J.R., Reidler, J.S., Sepulcre, J., Poulin, R., Buckner, R.L., 2010. Functional-anatomic fractionation of the brain’s default network. Neuron 65, 550–562.

Arbabshirani, M.R., Plis, S., Sui, J., Calhoun, V.D., 2017. Single subject prediction of brain disorders in neuroimaging: Promises and pitfalls. Neuroimage 145, 137–165.

Arthur, D., Vassilvitskii, S., 2007. k-means++: The advantages of careful seeding. Proceedings of the eighteenth annual ACM-SIAM symposium on Discrete algorithms. Society for Industrial and Applied Mathematics, pp. 1027–1035.

Baker, J.T., Holmes, A.J., Masters, G.A., Yeo, B.T., Krienen, F., Buckner, R.L., Ongur, D., 2014. Disruption of cortical association networks in schizophrenia and psychotic bipolar disorder. JAMA Psychiatry 71, 109–118.

Barttfeld, P., Uhrig, L., Sitt, J.D., Sigman, M., Jarraya, B., Dehaene, S., 2015. Signature of consciousness in the dynamics of resting-state brain activity. Proc Natl Acad Sci U S A 112, 887–892.

Beckmann, C.F., DeLuca, M., Devlin, J.T., Smith, S.M., 2005. Investigations into resting-state connectivity using independent component analysis. Philos Trans R Soc Lond B Biol Sci 360, 1001–1013.

Bell, A.J., Sejnowski, T.J., 1995. An information-maximization approach to blind separation and blind deconvolution. Neural Comput 7, 1129–1159.

Benjamini, Y., Hochberg, Y., 1995. Controlling the False Discovery Rate: A Practical and Powerful Approach to Multiple Testing. Journal of the Royal Statistical Society. Series B (Methodological) 57, 289–300.

Biswal, B.B., Mennes, M., Zuo, X.N., Gohel, S., Kelly, C., Smith, S.M., Beckmann, C.F., Adelstein, J.S., Buckner, R.L., Colcombe, S., Dogonowski, A.M., Ernst, M., Fair, D., Hampson, M., Hoptman, M.J., Hyde, J.S., Kiviniemi, V.J., Kotter, R., Li, S.J., Lin, C.P., Lowe, M.J., Mackay, C., Madden, D.J., Madsen, K.H., Margulies, D.S., Mayberg, H.S., McMahon, K., Monk, C.S., Mostofsky, S.H., Nagel, B.J., Pekar, J.J., Peltier, S.J., Petersen, S.E., Riedl, V., Rombouts, S.A., Rypma, B., Schlaggar, B.L., Schmidt, S., Seidler, R.D., Siegle, G.J., Sorg, C., Teng, G.J., Veijola, J., Villringer, A., Walter, M., Wang, L., Weng, X.C., Whitfield-Gabrieli, S., Williamson, P., Windischberger, C., Zang, Y.F., Zhang, H.Y., Castellanos, F.X., Milham, M.P., 2010. Toward discovery science of human brain function. Proc Natl Acad Sci U S A 107, 4734–4739.

Braga, R.M., Buckner, R.L., 2017. Parallel Interdigitated Distributed Networks within the Individual Estimated by Intrinsic Functional Connectivity. Neuron 95, 457–471 e455.

Bressler, D.W., Fortenbaugh, F.C., Robertson, L.C., Silver, M.A., 2013. Visual spatial attention enhances the amplitude of positive and negative fMRI responses to visual stimulation in an eccentricity-dependent manner. Vision Res 85, 104–112.

Buckner, R.L., Andrews-Hanna, J.R., Schacter, D.L., 2008. The brain’s default network: anatomy, function, and relevance to disease. Ann N Y Acad Sci 1124, 1–38.

Buckner, R.L., Krienen, F.M., Castellanos, A., Diaz, J.C., Yeo, B.T., 2011. The organization of the human cerebellum estimated by intrinsic functional connectivity. J Neurophysiol 106, 2322–2345.

Buckner, R.L., Sepulcre, J., Talukdar, T., Krienen, F.M., Liu, H., Hedden, T., Andrews-Hanna, J.R., Sperling, R.A., Johnson, K.A., 2009. Cortical hubs revealed by intrinsic functional connectivity: mapping, assessment of stability, and relation to Alzheimer’s disease. J Neurosci 29, 1860–1873.

Bullier, J., Hupe, J.M., James, A.C., Girard, P., 2001. The role of feedback connections in shaping the responses of visual cortical neurons. Prog Brain Res 134, 193–204.

Calhoun, V.D., Adali, T., 2012. Multisubject independent component analysis of fMRI: a decade of intrinsic networks, default mode, and neurodiagnostic discovery. IEEE Rev Biomed Eng 5, 60–73.

Calhoun, V.D., Adali, T., Pearlson, G.D., Pekar, J.J., 2001a. A method for making group inferences from functional MRI data using independent component analysis. Hum Brain Mapp 14, 140–151.

Calhoun, V.D., Adali, T., Pearlson, G.D., Pekar, J.J., 2001b. Spatial and temporal independent component analysis of functional MRI data containing a pair of task-related waveforms. Hum Brain Mapp 13, 43–53.

Calhoun, V.D., Eichele, T., Pearlson, G., 2009. Functional brain networks in schizophrenia: a review. Front Hum Neurosci 3, 17.

Calhoun, V.D., Kiehl, K.A., Pearlson, G.D., 2008. Modulation of temporally coherent brain networks estimated using ICA at rest and during cognitive tasks. Hum Brain Mapp 29, 828–838.

Calhoun, V.D., Miller, R., Pearlson, G., Adali, T., 2014. The chronnectome: time-varying connectivity networks as the next frontier in fMRI data discovery. Neuron 84, 262–274.

Cazorla, M., Kang, U.J., Kellendonk, C., 2015. Balancing the basal ganglia circuitry: a possible new role for dopamine D2 receptors in health and disease. Mov Disord 30, 895–903.

Chen, T., Cai, W., Ryali, S., Supekar, K., Menon, V., 2016. Distinct Global Brain Dynamics and Spatiotemporal Organization of the Salience Network. PLoS Biol 14, e1002469.

Correa, N., Adali, T., Calhoun, V.D., 2007. Performance of Blind Source Separation Algorithms for FMRI Analysis using a Group ICA Method. Magnetic resonance imaging 25, 684–694.

Correa, N., Adali, T., Li, Y.-O., Calhoun, V.D., 2005. Comparison of blind source separation algorithms for FMRI using a new Matlab toolbox: GIFT. Acoustics, Speech, and Signal Processing, 2005. Proceedings.(ICASSP’05). IEEE International Conference on. IEEE, pp. v/401–v/404 Vol. 405.

Damaraju, E., Allen, E.A., Belger, A., Ford, J.M., McEwen, S., Mathalon, D.H., Mueller, B.A., Pearlson, G.D., Potkin, S.G., Preda, A., Turner, J.A., Vaidya, J.G., van Erp, T.G., Calhoun, V.D., 2014. Dynamic functional connectivity analysis reveals transient states of dysconnectivity in schizophrenia. Neuroimage Clin 5, 298–308.

Damoiseaux, J.S., Rombouts, S.A.R.B., Barkhof, F., Scheltens, P., Stam, C.J., Smith, S.M., Beckmann, C.F., 2006. Consistent resting-state networks across healthy subjects. Proc Natl Acad Sci U S A 103, 13848–13853.

Du, Y., Fan, Y., 2013. Group information guided ICA for fMRI data analysis. Neuroimage 69, 157–197.

Du, Y., Pearlson, G.D., Liu, J., Sui, J., Yu, Q., He, H., Castro, E., Calhoun, V.D., 2015. A group ICA based framework for evaluating resting fMRI markers when disease categories are unclear: application to schizophrenia, bipolar, and schizoaffective disorders. Neuroimage 122, 272–280.

Erdeniz, B., Serin, E., Ibadi, Y., Tas, C., 2017. Decreased functional connectivity in schizophrenia: The relationship between social functioning, social cognition and graph theoretical network measures. Psychiatry Res Neuroimaging 270, 22–31.

Erhardt, E.B., Allen, E.A., Damaraju, E., Calhoun, V.D., 2011a. On network derivation, classification, and visualization: a response to Habeck and Moeller. Brain Connect 1, 105–110.

Erhardt, E.B., Rachakonda, S., Bedrick, E.J., Allen, E.A., Adali, T., Calhoun, V.D., 2011b. Comparison of multi-subject ICA methods for analysis of fMRI data. Hum Brain Mapp 32, 2075–2095.

Felleman, D.J., Van Essen, D.C., 1991. Distributed hierarchical processing in the primate cerebral cortex. Cereb Cortex 1, 1–47.

Fox, M.D., Corbetta, M., Snyder, A.Z., Vincent, J.L., Raichle, M.E., 2006. Spontaneous neuronal activity distinguishes human dorsal and ventral attention systems. Proc Natl Acad Sci U S A 103, 10046–10051.

Fox, M.D., Snyder, A.Z., Vincent, J.L., Corbetta, M., Van Essen, D.C., Raichle, M.E., 2005. The human brain is intrinsically organized into dynamic, anticorrelated functional networks. Proc Natl Acad Sci U S A 102, 9673–9678.

Fox, M.D., Zhang, D., Snyder, A.Z., Raichle, M.E., 2009. The global signal and observed anticorrelated resting state brain networks. J Neurophysiol 101, 3270–3283.

Franco, A.R., Pritchard, A., Calhoun, V.D., Mayer, A.R., 2009. Interrater and Intermethod Reliability of Default Mode Network Selection. Hum Brain Mapp 30, 2293–2303.

Friston, K.J., 1998. The disconnection hypothesis. Schizophr Res 30, 115–125.

Garrity, A.G., Pearlson, G.D., McKiernan, K., Lloyd, D., Kiehl, K.A., Calhoun, V.D., 2007. Aberrant “default mode” functional connectivity in schizophrenia. Am J Psychiatry 164, 450–457.

Gavrilescu, M., Rossell, S., Stuart, G.W., Shea, T.L., Innes-Brown, H., Henshall, K., McKay, C., Sergejew, A.A., Copolov, D., Egan, G.F., 2010. Reduced connectivity of the auditory cortex in patients with auditory hallucinations: a resting state functional magnetic resonance imaging study. Psychol Med 40, 1149–1158.

Greicius, M., 2008. Resting-state functional connectivity in neuropsychiatric disorders. Curr Opin Neurol 21, 424–430.

Greicius, M.D., Krasnow, B., Reiss, A.L., Menon, V., 2003. Functional connectivity in the resting brain: a network analysis of the default mode hypothesis. Proc Natl Acad Sci U S A 100, 253–258.

Guo, C.C., Kurth, F., Zhou, J., Mayer, E.A., Eickhoff, S.B., Kramer, J.H., Seeley, W.W., 2012. One-year test-retest reliability of intrinsic connectivity network fMRI in older adults. Neuroimage 61, 1471–1483.

Haralick, R.M., Shanmugam, K., Dinstein, I.H., 1973. Textural features for image classification. IEEE Transactions on systems, man, and cybernetics 3, 610–621.

Harrison, S.A., Tong, F., 2009. Decoding reveals the contents of visual working memory in early visual areas. Nature 458, 632–635.

Himberg, J., Hyvarinen, A., Esposito, F., 2004. Validating the independent components of neuroimaging time series via clustering and visualization. Neuroimage 22, 1214–1222.

Hupe, J.M., James, A.C., Girard, P., Lomber, S.G., Payne, B.R., Bullier, J., 2001. Feedback connections act on the early part of the responses in monkey visual cortex. J Neurophysiol 85, 134–145.

Hutchison, R.M., Womelsdorf, T., Allen, E.A., Bandettini, P.A., Calhoun, V.D., Corbetta, M., Della Penna, S., Duyn, J.H., Glover, G.H., Gonzalez-Castillo, J., Handwerker, D.A., Keilholz, S., Kiviniemi, V., Leopold, D.A., de Pasquale, F., Sporns, O., Walter, M., Chang, C., 2013. Dynamic functional connectivity: promise, issues, and interpretations. Neuroimage 80, 360–378.

Iraji, A., Benson, R.R., Welch, R.D., O’Neil, B.J., Woodard, J.L., Ayaz, S.I., Kulek, A., Mika, V., Medado, P., Soltanian-Zadeh, H., Liu, T., Haacke, E.M., Kou, Z., 2015. Resting State Functional Connectivity in Mild Traumatic Brain Injury at the Acute Stage: Independent Component, and Seed-Based Analyses. J Neurotrauma 32, 1031–1045.

Iraji, A., Calhoun, V.D., Wiseman, N.M., Davoodi-Bojd, E., Avanaki, M.R.N., Haacke, E.M., Kou, Z., 2016. The connectivity domain: Analyzing resting state fMRI data using feature-based data-driven and model-based methods. Neuroimage 134, 494–507.

Jafri, M.J., Pearlson, G.D., Stevens, M., Calhoun, V.D., 2008. A method for functional network connectivity among spatially independent resting-state components in schizophrenia. Neuroimage 39, 1666–1681.

Karahanoglu, F.I., Van De Ville, D., 2015. Transient brain activity disentangles fMRI resting-state dynamics in terms of spatially and temporally overlapping networks. Nat Commun 6, 7751.

Kelly, R.M., Strick, P.L., 2003. Cerebellar loops with motor cortex and prefrontal cortex of a nonhuman primate. J Neurosci 23, 8432–8444.

Kiviniemi, V., Vire, T., Remes, J., Elseoud, A.A., Starck, T., Tervonen, O., Nikkinen, J., 2011. A sliding time-window ICA reveals spatial variability of the default mode network in time. Brain Connect 1, 339–347.

Krienen, F.M., Buckner, R.L., 2009. Segregated fronto-cerebellar circuits revealed by intrinsic functional connectivity. Cereb Cortex 19, 2485–2497.

Lars, M., 2010. What are we missing here? Brain imaging evidence for higher cognitive functions in primary visual cortex V1. International Journal of Imaging Systems and Technology 20, 131–139.

Laumann, T.O., Gordon, E.M., Adeyemo, B., Snyder, A.Z., Joo, S.J., Chen, M.Y., Gilmore, A.W., McDermott, K.B., Nelson, S.M., Dosenbach, N.U., Schlaggar, B.L., Mumford, J.A., Poldrack, R.A., Petersen, S.E., 2015. Functional System and Areal Organization of a Highly Sampled Individual Human Brain. Neuron 87, 657–670.

Lee, T.W., Xue, S.W., 2018. Functional connectivity maps based on hippocampal and thalamic dynamics may account for the default-mode network. Eur J Neurosci 47, 388–398.

Leech, R., Sharp, D.J., 2014. The role of the posterior cingulate cortex in cognition and disease. Brain 137, 12–32.

Leonardi, N., Richiardi, J., Gschwind, M., Simioni, S., Annoni, J.M., Schluep, M., Vuilleumier, P., Van De Ville, D., 2013. Principal components of functional connectivity: a new approach to study dynamic brain connectivity during rest. Neuroimage 83, 937–950.

Liu, X., Duyn, J.H., 2013. Time-varying functional network information extracted from brief instances of spontaneous brain activity. Proc Natl Acad Sci U S A 110, 4392–4397.

Ma, S., Calhoun, V.D., Phlypo, R., Adali, T., 2014. Dynamic changes of spatial functional network connectivity in healthy individuals and schizophrenia patients using independent vector analysis. Neuroimage 90, 196–206.

Ma, S., Correa, N.M., Li, X.L., Eichele, T., Calhoun, V.D., Adali, T., 2011. Automatic identification of functional clusters in FMRI data using spatial dependence. IEEE Trans Biomed Eng 58, 3406–3417.

Markov, N.T., Misery, P., Falchier, A., Lamy, C., Vezoli, J., Quilodran, R., Gariel, M.A., Giroud, P., Ercsey-Ravasz, M., Pilaz, L.J., Huissoud, C., Barone, P., Dehay, C., Toroczkai, Z., Van Essen, D.C., Kennedy, H., Knoblauch, K., 2011. Weight consistency specifies regularities of macaque cortical networks. Cereb Cortex 21, 1254–1272.

Menon, V., 2011. Large-scale brain networks and psychopathology: a unifying triple network model. Trends Cogn Sci 15, 483–506.

Muckli, L., Petro, L.S., 2013. Network interactions: non-geniculate input to V1. Curr Opin Neurobiol 23, 195–201.

O’Reilly, J.X., Beckmann, C.F., Tomassini, V., Ramnani, N., Johansen-Berg, H., 2010. Distinct and overlapping functional zones in the cerebellum defined by resting state functional connectivity. Cereb Cortex 20, 953–965.

Power, J.D., Barnes, K.A., Snyder, A.Z., Schlaggar, B.L., Petersen, S.E., 2012. Spurious but systematic correlations in functional connectivity MRI networks arise from subject motion. Neuroimage 59, 2142–2154.

Preti, M.G., Bolton, T.A., Van De Ville, D., 2017. The dynamic functional connectome: State-of-the-art and perspectives. Neuroimage 160, 41–54.

Preti, M.G., Van De Ville, D., 2017. Dynamics of functional connectivity at high spatial resolution reveal long-range interactions and fine-scale organization. Sci Rep 7, 12773.

Roelfsema, P.R., de Lange, F.P., 2016. Early Visual Cortex as a Multiscale Cognitive Blackboard. Annu Rev Vis Sci 2, 131–151.

Sakoglu, U., Pearlson, G.D., Kiehl, K.A., Wang, Y.M., Michael, A.M., Calhoun, V.D., 2010. A method for evaluating dynamic functional network connectivity and task-modulation: application to schizophrenia. MAGMA 23, 351–366.

Seeley, W.W., Crawford, R.K., Zhou, J., Miller, B.L., Greicius, M.D., 2009. Neurodegenerative diseases target large-scale human brain networks. Neuron 62, 42–52.

Shehzad, Z., Kelly, A.M., Reiss, P.T., Gee, D.G., Gotimer, K., Uddin, L.Q., Lee, S.H., Margulies, D.S., Roy, A.K., Biswal, B.B., Petkova, E., Castellanos, F.X., Milham, M.P., 2009. The resting brain: unconstrained yet reliable. Cereb Cortex 19, 2209–2229.

Shine, J.M., Koyejo, O., Poldrack, R.A., 2016. Temporal metastates are associated with differential patterns of time-resolved connectivity, network topology, and attention. Proc Natl Acad Sci U S A 113, 9888–9891.

Shirer, W.R., Ryali, S., Rykhlevskaia, E., Menon, V., Greicius, M.D., 2012. Decoding subject-driven cognitive states with whole-brain connectivity patterns. Cereb Cortex 22, 158–165.

Sincich, L.C., Park, K.F., Wohlgemuth, M.J., Horton, J.C., 2004. Bypassing V1: a direct geniculate input to area MT. Nat Neurosci 7, 1123–1128.

Skudlarski, P., Jagannathan, K., Anderson, K., Stevens, M.C., Calhoun, V.D., Skudlarska, B.A., Pearlson, G., 2010. Brain connectivity is not only lower but different in schizophrenia: a combined anatomical and functional approach. Biol Psychiatry 68, 61–69.

Smith, S.M., Fox, P.T., Miller, K.L., Glahn, D.C., Fox, P.M., Mackay, C.E., Filippini, N., Watkins, K.E., Toro, R., Laird, A.R., Beckmann, C.F., 2009. Correspondence of the brain’s functional architecture during activation and rest. Proc Natl Acad Sci U S A 106, 13040–13045.

Smith, S.M., Miller, K.L., Moeller, S., Xu, J., Auerbach, E.J., Woolrich, M.W., Beckmann, C.F., Jenkinson, M., Andersson, J., Glasser, M.F., Van Essen, D.C., Feinberg, D.A., Yacoub, E.S., Ugurbil, K., 2012. Temporally-independent functional modes of spontaneous brain activity. Proc Natl Acad Sci U S A 109, 3131–3136.

Sorg, C., Riedl, V., Muhlau, M., Calhoun, V.D., Eichele, T., Laer, L., Drzezga, A., Forstl, H., Kurz, A., Zimmer, C., Wohlschlager, A.M., 2007. Selective changes of resting-state networks in individuals at risk for Alzheimer’s disease. Proc Natl Acad Sci U S A 104, 18760–18765.

Stoodley, C.J., Schmahmann, J.D., 2009. Functional topography in the human cerebellum: a meta-analysis of neuroimaging studies. Neuroimage 44, 489–501.

Strick, P.L., Dum, R.P., Fiez, J.A., 2009. Cerebellum and nonmotor function. Annu Rev Neurosci 32, 413–434.

Tagliazucchi, E., Balenzuela, P., Fraiman, D., Chialvo, D.R., 2012. Criticality in large-scale brain FMRI dynamics unveiled by a novel point process analysis. Front Physiol 3, 15.

Trapp, C., Vakamudi, K., Posse, S., 2018. On the detection of high frequency correlations in resting state fMRI. Neuroimage 164, 202–213.

Uddin, L.Q., Kelly, A.M., Biswal, B.B., Castellanos, F.X., Milham, M.P., 2009. Functional connectivity of default mode network components: correlation, anticorrelation, and causality. Hum Brain Mapp 30, 625–637.

van de Ven, V., Jacobs, C., Sack, A.T., 2012. Topographic contribution of early visual cortex to short-term memory consolidation: a transcranial magnetic stimulation study. J Neurosci 32, 4–11.

van den Heuvel, M.P., Mandl, R.C., Kahn, R.S., Hulshoff Pol, H.E., 2009. Functionally linked resting-state networks reflect the underlying structural connectivity architecture of the human brain. Hum Brain Mapp 30, 3127–3141.

Van Dijk, K.R., Hedden, T., Venkataraman, A., Evans, K.C., Lazar, S.W., Buckner, R.L., 2010. Intrinsic functional connectivity as a tool for human connectomics: theory, properties, and optimization. J Neurophysiol 103, 297–321.

van Erp, T.G., Preda, A., Turner, J.A., Callahan, S., Calhoun, V.D., Bustillo, J.R., Lim, K.O., Mueller, B., Brown, G.G., Vaidya, J.G., McEwen, S., Belger, A., Voyvodic, J., Mathalon, D.H., Nguyen, D., Ford, J.M., Potkin, S.G., 2015. Neuropsychological profile in adult schizophrenia measured with the CMINDS. Psychiatry Res 230, 826–834.

Vercammen, A., Knegtering, H., den Boer, J.A., Liemburg, E.J., Aleman, A., 2010. Auditory hallucinations in schizophrenia are associated with reduced functional connectivity of the temporo-parietal area. Biol Psychiatry 67, 912–918.

Vidaurre, D., Smith, S.M., Woolrich, M.W., 2017. Brain network dynamics are hierarchically organized in time. Proc Natl Acad Sci U S A 114, 12827–12832.

Wang, X., Xu, M., Song, Y., Li, X., Zhen, Z., Yang, Z., Liu, J., 2014. The network property of the thalamus in the default mode network is correlated with trait mindfulness. Neuroscience 278, 291–301.

Yaesoubi, M., Adali, T., Calhoun, V.D., 2018. A window-less approach for capturing time-varying connectivity in fMRI data reveals the presence of states with variable rates of change. Hum Brain Mapp 39, 1626–1636.

Yaesoubi, M., Miller, R.L., Calhoun, V.D., 2017. Time-varying spectral power of resting-state fMRI networks reveal cross-frequency dependence in dynamic connectivity. PLoS One 12, e0171647.

Yeo, B.T., Krienen, F.M., Sepulcre, J., Sabuncu, M.R., Lashkari, D., Hollinshead, M., Roffman, J.L., Smoller, J.W., Zollei, L., Polimeni, J.R., Fischl, B., Liu, H., Buckner, R.L., 2011. The organization of the human cerebral cortex estimated by intrinsic functional connectivity. J Neurophysiol 106, 1125–1165.

Zeng, L.L., Wang, H., Hu, P., Yang, B., Pu, W., Shen, H., Chen, X., Liu, Z., Yin, H., Tan, Q., Wang, K., Hu, D., 2018. Multi-Site Diagnostic Classification of Schizophrenia Using Discriminant Deep Learning with Functional Connectivity MRI. EBioMedicine 30, 74–85.

Zuo, X.N., Kelly, C., Adelstein, J.S., Klein, D.F., Castellanos, F.X., Milham, M.P., 2010. Reliable intrinsic connectivity networks: test-retest evaluation using ICA and dual regression approach. Neuroimage 49, 2163–2177.

